# Defining the cell surface proteomic landscape of multiple myeloma reveals immunotherapeutic strategies and biomarkers of drug resistance

**DOI:** 10.1101/2021.01.17.427038

**Authors:** Ian D. Ferguson, Bonell Patiño Escobar, Sami T. Tuomivaara, Yu-Hsiu T. Lin, Matthew A. Nix, Kevin K. Leung, Martina Hale, Priya Choudhry, Antonia Lopez-Girona, Emilio Ramos, Sandy W. Wong, Jeffrey L. Wolf, Thomas G. Martin, Nina Shah, Scott Vandenberg, Sonam Prakash, Lenka Besse, Christoph Driessen, James A. Wells, Arun P. Wiita

## Abstract

The myeloma cell surface proteome (“surfaceome”) not only determines tumor interaction with the microenvironment but serves as an emerging arena for therapeutic development. Here, we use glycoprotein capture proteomics to first define surface markers most-enriched on myeloma when compared to B-cell malignancy models, revealing unexpected biological signatures unique to malignant plasma cells. We next integrate our proteomic dataset with existing transcriptome databases, nominating CCR10 and TXNDC11 as possible monotherapeutic targets and CD48 as a promising co-target for increasing avidity of BCMA-directed cellular therapies. We further identify potential biomarkers of resistance to both proteasome inhibitors and lenalidomide including changes in CD53, EVI2B, CD10, and CD33. Comparison of short-term treatment with chronic resistance delineates large differences in surface proteome profile under each type of drug exposure. Finally, we develop a miniaturized version of the surface proteomics protocol and present the first surface proteomic profile of a primary myeloma patient plasma cell sample. Our dataset provides a unique resource to advance the biological, therapeutic, and diagnostic understanding of myeloma.

## INTRODUCTION

The composition of the tumor cell surface plays a central role in determining cancer’s interaction with the local microenvironment. Over the past several years, targeting surface proteins expressed on tumor cells has also rapidly emerged as one of the most exciting frontiers for treating cancer. This strategy is particularly relevant in the case of the plasma cell malignancy multiple myeloma. Antibody-based therapeutics targeting CD38, SLAMF7 and BCMA have now been FDA-approved, and engineered cellular therapies targeting BCMA have shown highly promising clinical data. Furthermore, identifying and quantifying cell surface markers play a critical role in the diagnosis and monitoring of all hematologic malignancies, including myeloma.

One of the notable clinical features of myeloma is that despite the recent advent of many effective therapies, there is still no known cure for this disease. Resistance to current small molecule therapeutics, particularly proteasome inhibitors such as bortezomib and carfilzomib, and immunomodulatory drugs such as lenalidomide, is a widespread conundrum. Characterizing surface proteomic changes in these contexts may reveal new strategies to diagnose and specifically treat drug-resistant disease.

Despite the importance of the cell surface to diagnosis, therapy, and biology of myeloma, much remains unknown. While extensive RNA-seq datasets are available on both myeloma primary samples (such as the Multiple Myeloma Research Foundation CoMMpass study (research.themmrf.org)) and cell lines (keatslab.org and Cancer Cell Line Encyclopedia (1)), it is well-known that transcript-level expression is only modestly correlated with surface protein expression (2–4). This lack of predictive power is related to two main features. First, even for proteins exclusively present at the plasma membrane, transcript-only quantification cannot capture alterations in translational regulation and protein trafficking that ultimately govern surface expression. Second, proteins expressed at the cell surface can also have significant pools of either intracellular or secreted forms as well; RNA-seq cannot distinguish these components. Therefore, these datasets can at best be considered partially predictive of the cell surface proteome.

Other studies have used flow cytometry, or, more recently, mass cytometry/CyTOF (5), to profile myeloma tumor cells in the context of inter-patient heterogeneity (6), response to therapy (7), or drug resistance (8). However, these assays are typically restricted to monitoring a maximum of ~50 surface proteins that are already very well-characterized and have high-quality antibodies available. Even a recent largescale survey of normal human B-cells by CyTOF, screening an exhaustive catalog of 351 metal-conjugated antibodies, only identified 98 expressed surface proteins (9). It is estimated that most human cells express >2000 unique proteins on their surface (10). Therefore, these widely-used approaches can only begin to outline the overall profile of the myeloma cell surface.

To overcome these limitations, here we use a relatively unbiased approach, glycoprotein cell surface capture (CSC) (11), to directly quantify hundreds of proteins localized to the surface of myeloma tumor cells. We specifically identify new potential immunotherapy strategies in myeloma and new biomarkers of small molecule resistance and response to acute treatment. Finally, to streamline the surface proteomics methodology we develop a miniaturized CSC approach and apply this approach to primary patient myeloma. Uncovering the surface landscape of malignant plasma cells serves as a significant resource to the myeloma community.

## RESULTS

### Determining the malignant plasma cell surface landscape

We first utilized the CSC approach to oxidize, covalently biotinylate, and then isolate N-linked glycoproteins from four multiple myeloma cell lines (KMS-12PE, AMO-1, RPMI-8226, L363) (**Fig. 1A**). Labeling was performed on live cells to generate an initial surface proteomic landscape of malignant plasma cells. In our initial analysis, using the label-free quantification approach in MaxQuant (12), we cumulatively quantified 1245 proteins annotated as membrane bound in Uniprot across all cell lines (ranging from 715-1069 in individual cell lines, with a minimum of two peptides per protein), with a common intersection of 562 proteins (**Fig. 1B**). To maximize number of captured proteins, we used glycoprotein biotinylation with on-bead trypsinization (**Fig. 1A**); however, this method is likely to also spuriously elute peptides deriving from background intracellular proteins. Filtering our data with a recently-described set of the best-validated plasma membrane proteins (10), 530 of these quantified proteins (305-436 per cell line) appear localized to the cell surface with high confidence (**Supp. Fig. 1A**). Notably, across these lines we detected almost all of the major immunotherapy targets in myeloma, as well as canonical flow cytometry markers for plasma cells (7, 13): BCMA, CD138/SDC1, CD38, CD56, SLAMF7/CS-1, CD46, Integrin-b7 (ITGB7), CD74/HLA-DR, TACI, CD48/SLAMF2 and LY9/CD229 (**Fig. 1C**; **Supplementary Dataset 1**).

**Figure 1.**
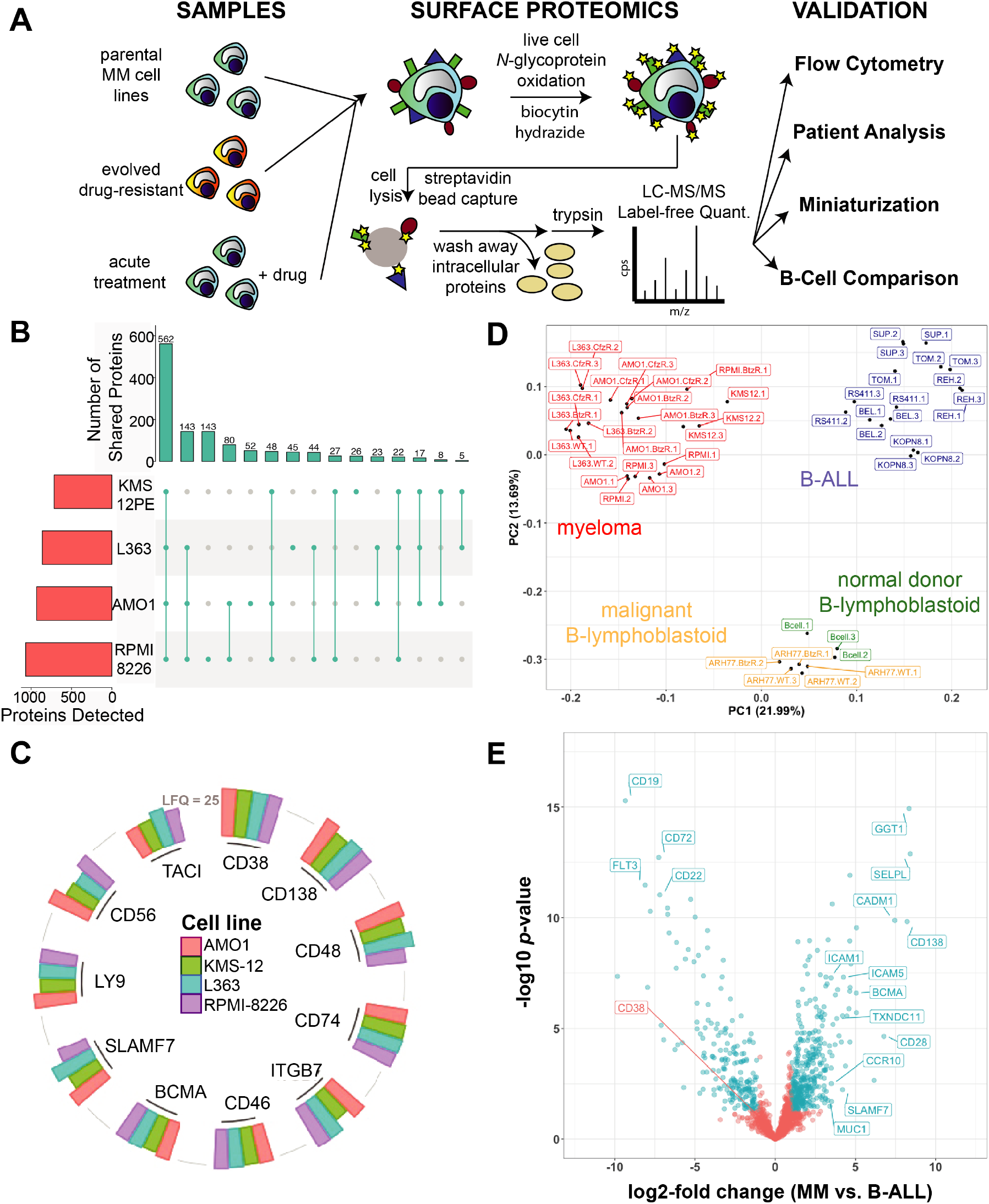
Initial elucidation of the myeloma plasma cell “surfaceome”. **A**. Overall schematic of surface proteomic investigations in this study. This includes a description of the modified cell surface capture (CSC) methodology used, with biotinylated proteins identified after on-bead trypsinization. **B.** Upset plot shows high degree of overlap in identified glycoproteins across the four evaluated myeloma cell lines. Data included if identified with two peptides in at least one of three biological replicate per cell line. **C.** Common myeloma diagnostic markers and immunotherapeutic targets were identified by cell surface proteomics in all four evaluated cell lines. Height of column indicates labelfree quantification (LFQ) intensity from MaxQuant, averaged across replicate samples. A threshold of LFQ = 25 is indicated by grey line. **D**. Principal Component Analysis (PCA) illustrates the differential cell surface landscape of myeloma plasma cells versus B-lymphoblastoid cells and B-cell acute lymphoblastic leukemia cell lines. BtzR and CfzR are bortezomib and carfilzomib resistant cells *(n* = 2 or 3 biological replicates per cell line, except for RPMI-BtzR). **E**. Volcano plot comparing glycoprotein LFQ intensity of four myeloma cell lines to eight B-ALL cell lines. Significantly changed proteins labeled in blue (log2-fold change >|1|; *p* < 0.05).

While these positive identifications are certainly encouraging for the utility of this dataset, we note this technology still cannot detect all cell surface proteins. For example, we did not detect GPRC5D, a G-protein coupled receptor (GPCR) recently proposed as a specific immunotherapy target in myeloma (14). In general, GPCRs and other multi-pass membrane proteins tend to be under-represented in CSC (4); we noted the same effect in our dataset (**Supp. Fig. 1B**). Furthermore, as in essentially all proteomic methods, proteins present at low copy number on the cell surface are less likely to be detected on the mass spectrometer (15). In addition, ~10-20% of surface proteins likely do not feature any *N*-linked glycosylation (16), and therefore will not be detectable via CSC. Despite these limitations, we propose that this initial landscape of the myeloma cell surface provides a unique resource for downstream investigation.

As myeloma is a malignancy of plasma cells, we were curious what surface markers particularly distinguish myeloma from B-cells earlier in the developmental trajectory. We therefore compared our myeloma surfaceome to our earlier dataset (17) including six B-cell acute lymphoblastic leukemia (B-ALL) lines derived from early (pre- or pro-) B-cell developmental stages. We further compared the myeloma surface profile to two B-lymphoblastoid cell lines (EBV-immortalized normal cord blood donor B-cells and ARH-77), derived from differentiated, late-stage B-cells (18). By Principal Component Analysis (PCA) we were encouraged to find that lines representing each cell type (plasma cells, early B-cell, late B-cell) clustered separately, consistent with a relatively unique surface signature for each cell type (**Fig. 1D**).

We next identified the specific markers that most distinguish myeloma plasma cells versus these other B-cell types (**Fig. 1D**; **Supp. Fig. 1C**). Giving confidence in this analysis, many canonical markers were among the strongest hits including BCMA, CD138/SDC1, and CD28 for plasma cells and CD19, CD22, and CD72 for B-ALL cells, respectively (**Fig. 1D**). Intriguingly, by intensity of mass spectrometry signal, CD38 was not significantly different between these cell types, underscoring possible clinical relevance of anti-CD38 monoclonal antibody therapy in B-ALL (19). We were surprised to find the three proteins that most-distinguished myeloma plasma cells from early-stage malignant B-cells, based on fold-change and *p*-value, were not canonical markers at all: gamma-glutamyl transferase 1 *(GGT1),* selectin-P ligand *(SELPLG),* and cell adhesion molecule 1 *(CADM1)* (**Fig. 1E**). Comparison of myeloma cells to the two B-lymphoblastoid cell lines also showed CD138/SDC1 and CD28 as characterizing myeloma cells, while CD19, CD22, and CD20 characterized late-stage B-cells. SLAMF6/CD352 and SLAMF1/CD150 most strongly differentiated B-lymphoblastoid cell lines, while adenylate cyclase 3 *(ADCY3),* Podocalyxin-like *(PODXL),* and Integrin-a3/ITA3/CD49c *(ITGA3)* were the most strongly-differentiating on myeloma cells (**Supp. Fig. 1C**). These proteins may carry previously unexplored diagnostic or biological relevance to myeloma or broader plasma cell biology.

In parallel with our surface proteomic data, we also obtained RNA-seq data for all of the analyzed cell lines. We overall found a modest quantitative correlation (Pearson *R* = 0.54; *p* < 2.2e-16) between the transcriptome and surface proteome of myeloma and BALL cells (**Supp. Fig. 1D**), consistent with prior studies (17, 20). We did find a set of proteins, including CD99, GGT1 and TOR1AIP2, with transcript level discordant with surface protein, suggestive of possible post-transcriptional regulation (**Supp. Fig. 1D**). Taken together, this initial profiling underscores the unique surface phenotype of malignant plasma cells and unexpected markers distinguishing them from closely related cellular models.

### Identifying new targets for myeloma antigen-specific immunotherapies

We next turned our attention to possible new immunotherapeutic strategies revealed by our surface profiling. We first integrated our proteomic data with publicly-available transcriptome datasets to create a ranking system for possible single-antigen immunotherapy targets, based on five criteria related to surface abundance and specificity for plasma cells (**Fig. 2A**; **Supplementary Table 1**). Emphasizing the validity of this strategy, four of the top six targets by our ranking either already have FDA-approved therapeutics or are being clinically investigated in myeloma: BCMA *(TNRFSF17* gene), TACI *(TNFRSF13B),* Integrin beta-7 *(ITGB7),* and CS-1/SLAMF7 *(SLAMF7)* (**Fig. 2A**). CD38, CD138/SDC1, LY9/CD229, CD48, and GPRC5D also ranked highly (**Supplementary Dataset 2**). Based on these results, we probed other high-scoring proteins found in our proteomic data that, to our knowledge, have not yet been explored as therapeutic targets in myeloma. Here we offer two particularly intriguing examples: CCR10 and TXNDC11. CCR10 is a chemokine receptor previously shown to be expressed on plasma cells and thought to relate to homing to resident tissues (21). We found this gene to be robustly expressed on myeloma plasma cells per the CCLE but with minimal expression on other tumor cell lines (**Supp. Fig. 2A**). Data from GTex also suggest low, though not absent, mRNA expression on non-hematopoietic tissues, while data included in the Human Blood Atlas suggest markedly higher mRNA expression on plasmablasts than other hematopoietic cells, with the exception of some T-cell subtypes, including T regulatory cells, consistent with prior literature (22) (**Supp. Fig. 2A**). By flow cytometry we verified markedly increased CCR10 expression on myeloma cell lines compared to B-cell malignancy lines, of the same magnitude as BCMA relative increase, and also confirmed CCR10 expression on myeloma patient tumor cells (**Supp. Fig. 2B**). Together, these features suggest CCR10 as a promising immunotherapeutic target in myeloma.

**Figure 2.**
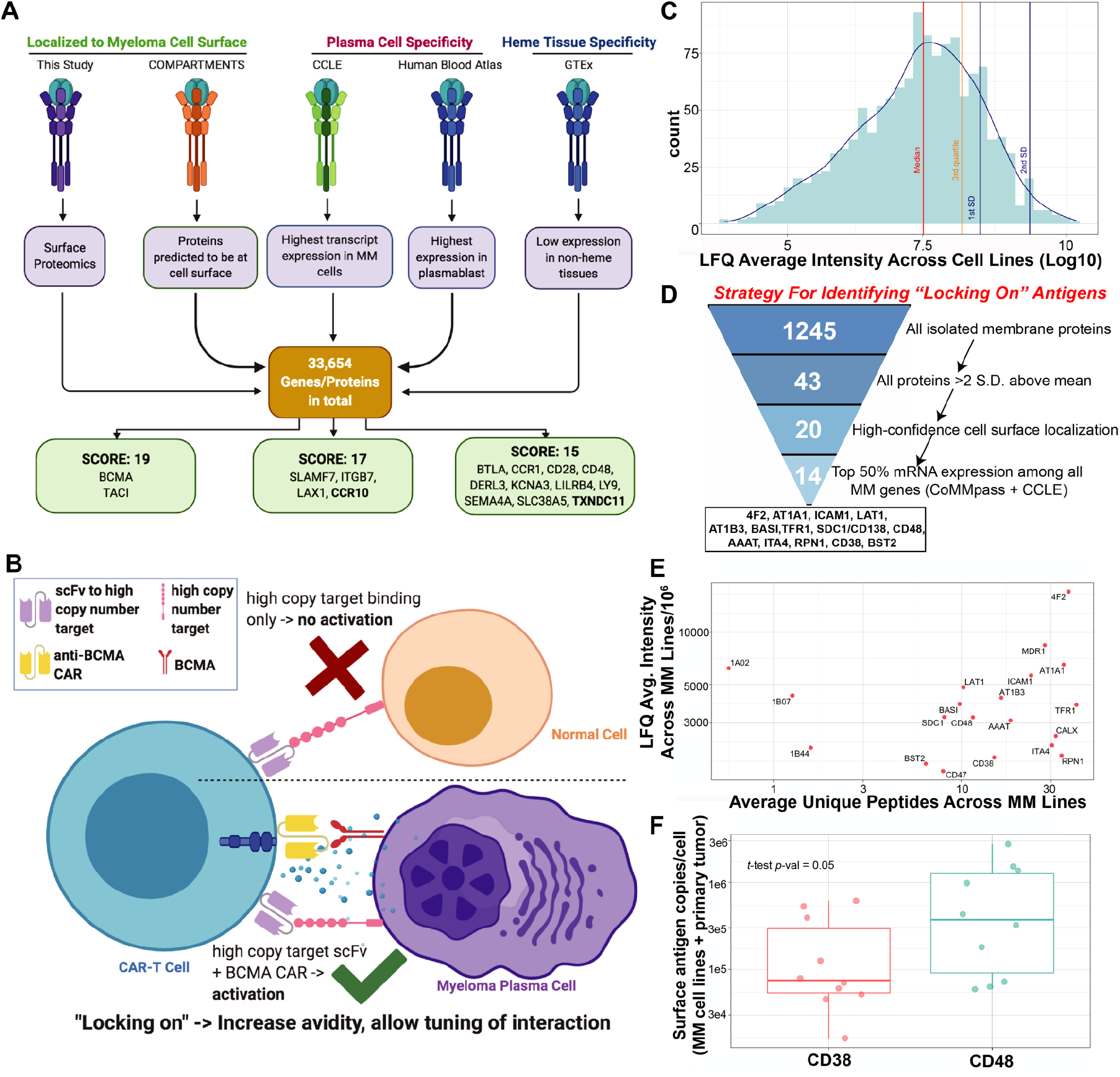
Immunotherapeutically targeting the myeloma cell surfaceome. **A.** Outline of a five-criteria scoring strategy, integrating surface proteomics data here with publicly-available mRNA transcriptome data, to propose new targets for possible antigen-specific immunotherapies in myeloma (see Methods for details). We specifically point out surface proteins with the highest scores among the total 33,654 analyzed (see **Supplementary Table 1** for scoring rubric; maximum score = 19). **B**. Schematic illustrating the concept of using a high-abundance, “locking on” antigen to tune CAR-T activation via increased avidity. **C.** Distribution of mass spectrometric intensity (label-free quantification (LFQ) from MaxQuant) of all glycoproteins quantified in our proteomic data; intensity is averaged across all four analyzed cell lines. **D.** Bioinformatic strategy to nominate possible high-abundance “locking on” antigens. **E.** Plot displaying the average LFQ intensity across cell lines of proteins >2 S.D. above the mean (C) versus average unique peptides identified in each line. **F.** Absolute quantification by flow cytometry for CD38 and CD48 antigen density across 3 myeloma cell lines (MM.1S, OPM-2, AMO-1) and CD138+/CD19-myeloma tumor cells from 5 primary patient bone marrow specimens. Box plot shows median and interquartile range of surface antigen density. *t*-value by *t-* test. (A) and (B) created with BioRender.

While TXNDC11 is generally poorly characterized, its sparse literature revolves around its role in endoplasmic reticulum (ER) associated protein degradation (23, 24). However, proteins with known ER localization may also have surface components; for example, the ER-resident HSP70 isoform BiP/GRP78 can be found at the cell surface in myeloma and serve as an immunotherapy target (25). Furthermore, immunofluorescence data in the Human Protein Atlas (26), as well as bioinformatic prediction from COMPARTMENTS (27), both place the primary location of TXNDC11 at the plasma membrane, not the ER. Similar to CCR10, while some non-hematopoietic tissues express TXNDC11, expression appears to be highly enhanced on plasma cells (**Supp. Fig. 2C**). In addition, per the Cancer Dependency Map (28), myeloma plasma cells appear somewhat genetically dependent on TXNDC11 for proliferation (**Supp. Fig. 2C**), potentially reducing probability of tumor resistance via antigen loss under therapeutic pressure. Unfortunately, we could not validate any currently available antibodies against TXNDC11 as suitable for flow cytometry (not shown). Therefore, we cannot definitively confirm that TXNDC11 spans the plasma membrane, nor rule out that TXNDC11 is identified in our dataset due to spurious background labeling of intracellular glycoproteins. Future studies will probe the relevance of this potential target. While we highlight these two proteins, additional targets from our integrative analysis may also warrant further follow-up in myeloma therapy.

### Characterizing the most abundant myeloma surface proteins for potential cotargeting

We next reasoned that identifying the most highly-expressed proteins on the surface of plasma cells may be advantageous for certain therapeutic designs. We illustrate one potential approach in **Fig. 2B**. In this embodiment, an scFv with no intracellular signaling domains (i.e. not a full CAR construct), specifically designed for “locking on” against a high abundance tumor antigen, would be highly expressed on the surface of the engineered T-cell. However, the T-cell would only be activated after engagement of an anti-BCMA (or other highly myeloma-specific antigen) scFv incorporated into a CAR. This approach would be designed to specifically increase the avidity of CAR-T binding to tumor. We envision that this approach, by increasing T-cell dwell time on tumor, could potentially enable CAR-T killing at lower antigen densities, which recent studies have suggested may be an important determinant of efficacy (29, 30). Or, alternatively, this approach may allow for reducing CAR signaling strength to minimize exhaustion (31) while still ultimately achieving the same degree of tumor killing, given more prolonged association between T-cells and target cells.

In this approach, the highly abundant tumor antigen for “locking on” would not necessarily have to be extremely specific for plasma cells, as no killing would occur if BCMA (or other targeted myeloma-specific antigen) were not present. We therefore interrogated our data to identify proteins that were among 1) most highly abundant on myeloma plasma cells, 2) confidently localized to the cell surface, 3) highly expressed at transcript level in myeloma patients per the CoMMpass database; 4) highly expressed on myeloma cell lines per CCLE data (see Methods for details). This analysis identified 14 potential “locking on” protein candidates (**Fig. 2D**).

We additionally probed biological signatures of surface proteins >2 S.D. above the LFQ mean (**Fig. 2E**). 4F2 (encoded by *SLC3A2)* and LAT1 *(SLC7A5),* which together comprise the heterodimeric large neutral amino acid transporter CD98, as well as the neutral amino acid transporter AAAT *(SLC1A5),* appear to be the most abundant proteins on the myeloma cell surface. This observation is potentially consistent with the critical role of protein synthesis in plasma cell biology. Other high-abundance plasma membrane proteins govern homeostatic mechanisms common across most human cells (AT1A1, AT1B3, TFR1/CD71). However, several of the other most highly abundant surface proteins carry specific functions in cellular signaling, either in myeloma plasma cells or hematopoietic cells more generally, including CD138/SDC1, CD47, CD38, ICAM1/CD54, ITA4/CD49d, and CD48/SLAMF2.

Integrating these two analyses, we specifically focused on two targets for further evaluation as “locking on” antigens: CD38 and CD48. We chose these proteins as others nominated in our analyses of **Fig. 2D-E** were expressed on at least some non-hematopoeitic tissues per GTex (**Supp. Fig. 3A**), potentially leading to misdirection of CAR-T’s during the locking-on phase by homing to irrelevant body sites. We particularly note that some of these proteins, most prominently CD38, would not be identified as highly abundant on the cell surface purely based on mRNA-seq analysis, illustrating the utility of proteomic studies in this application (**Supp. Fig. 3B**). CD38, a cell surface ectoenzyme, is well-known in myeloma as the target of the monoclonal antibodies (mAbs) daratumumab and isatuximab (32). While CD38 has been proposed as a standalone CAR-T target in myeloma, concern remains about toxicity given CD38 expression on many other hematopoietic cells and increased potency of CAR-Ts at lower antigen densities when compared to mAbs (13). CD48/SLAMF2, a CD2 ligand, has been proposed as a therapeutic target in myeloma, with prior studies confirming that >90% of myeloma primary samples tested expressed CD48 (33). Notably, CD48, but not CD38, shows greater transcript expression on myeloma cell lines than any other tumor cell type in the CCLE (**Supp. Fig. 3C**). However, concerns about toxicity still arise with CD48 monotherapy, given relatively high expression across other hematopoietic cells (33). Therefore, using either of these antigens as “locking on” handles, which do not activate T-cells when engaged, may be the most effective way to deploy them in myeloma cellular therapy.

To evaluate which of these proteins is more highly expressed at the myeloma cell surface, and thus best-positioned to increase T-cell avidity, we used a calibrated fluorescence strategy by flow cytometry to measure absolute antigen density of both CD38 and CD48 expression. We evaluated myeloma cell lines (MM.1S, OPM-2, AMO-1) and primary bone marrow aspirate samples from 5 relapsed/refractory myeloma patients, selected on CD19-/CD138+ plasma cells. We found that both antigens were expressed at high copy number, consistent with our proteomic findings, but CD48 showed the higher antigen density (range: 59,307-2,769,932 copies/cell vs. 16,251-613,422 for CD38) *(p* = 0.05) (**Fig. 2F**; **Supp. Fig. 3D**). We thus conclude that CD48 may be a particularly strong candidate for a potential “locking on” strategy to enhance the efficacy of BCMA-directed CAR-T’s.

### The myeloma surface proteome is remodeled in the context of proteasome inhibitor resistance

Having characterized the surface proteome of plasma cells at baseline, we next proposed that cataloging alterations in the context of proteasome inhibitor (PI) resistance may be relevant for diagnosing this condition, identifying new biological strategies to overcome resistance, or developing immunotherapies that selectively eliminate resistant disease. To this end, we performed cell surface proteomic profiling of AMO1, L363, and RPMI-8226 myeloma cell lines previously described to be *in vitro-evolved* for resistance to either carfilzomib (CfzR) or bortezomib (BtzR) (34) (**Fig. 3A**; **Supp Fig. 4A-B**, **Supplementary Dataset 3**). Serving as a positive control, the drug efflux pump MDR1 *(ABCB1)* was by far the most increased surface protein in CfzR cells, consistent with our prior results from whole cell shotgun proteomics (35, 36) (**Supp. Fig. 4C**). Aggregating BtzR vs. wild-type data, though, we saw no change in MDR1 (**Supp. Fig. 4D**), consistent with prior findings (35). Instead, across both CfzR and BtzR comparisons we found a signature whereby CD50, CD361/EVI2B, CD53, and Integrin-b7 (ITGB7) were commonly decreased while CD10 and CD151 were increased versus parental. Compared to RNA-seq (**Supplementary Dataset 4**), several genes showed evidence of possible post-transcriptional regulation; CD151, for example, showed >2.5-fold increased surface protein but essentially no transcript change in AMO1-BtzR cells (**Supp. Fig. 4E**). We did not identify significant upregulation of any current myeloma immunotherapy target proteins (**Supplementary Dataset 3**). Furthermore, with the possible exception of CD10, we did not identify signatures suggestive of de-differentiation to a more B-cell-like surface protein profile (37).

**Figure 3.**
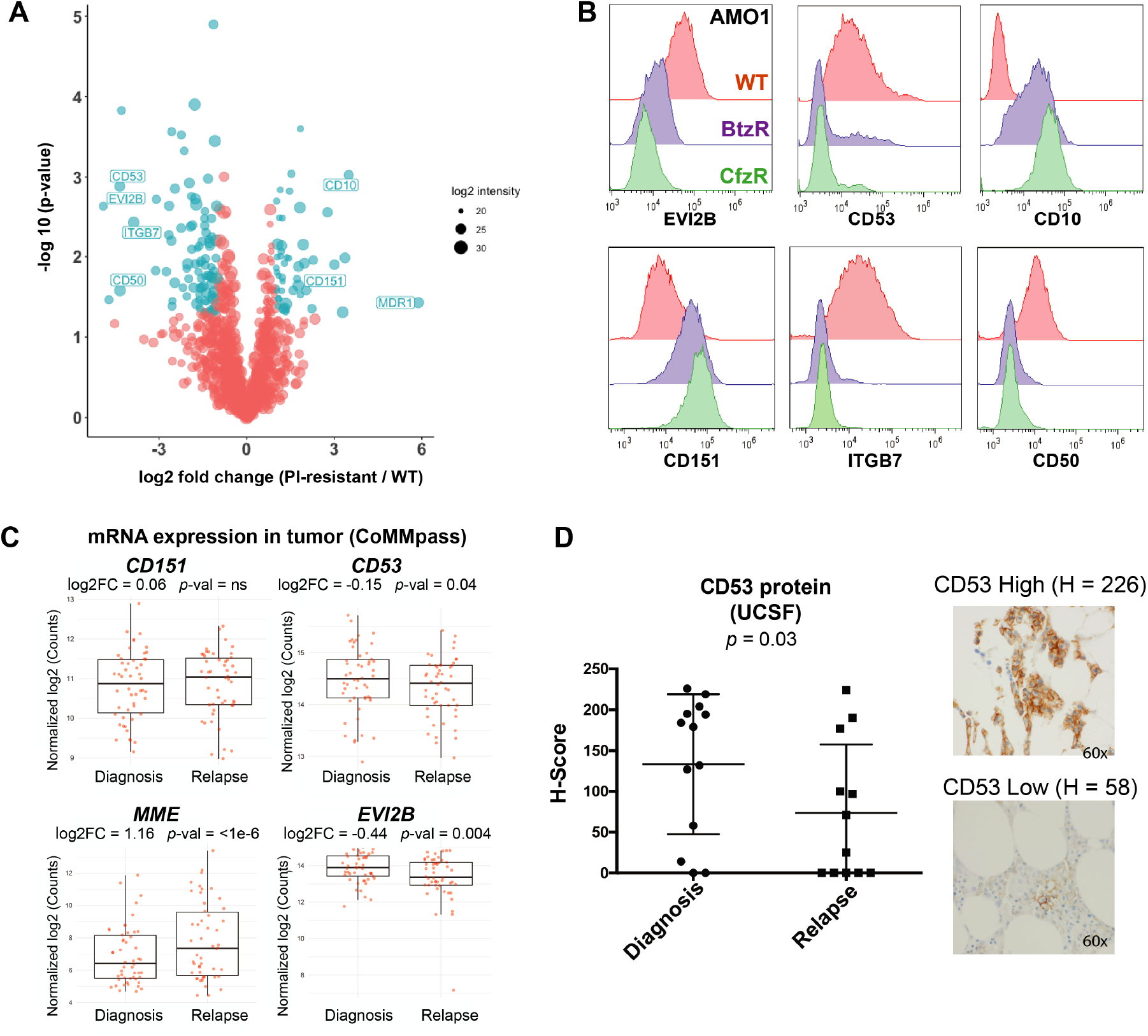
Defining a myeloma surface signature of proteasome inhibitor resistance. **A.** Cell surface proteomics was performed on evolved bortezomib-resistant (BtzR) and carfilzomib resistant (CfzR) myeloma cell lines (AMO-1 BtzR *(n* = 3); AMO1 CfzR *(n* = 3), L363 BtzR (*n* = 2), L363 CfzR (*n* = 3), RPMI-8226 BtzR (*n* = 1)) and aggregated in comparison to wild-type cell lines (AMO1 (*n* = 3), L363 (*n* = 2), RPMI-8226 (*n* = 1)). Significantly changed proteins in PI-resistant lines shown in blue (log2-fold change >|1|; *p* < 0.05). **B.** Validation by flow cytometry of most-changed surface proteins in AMO-1 cells. **C.** mRNA data in the MMRF CoMMpass database (Release IA14) from paired diagnosis and first-relapse tumor cells (*n* = 50), where all patients had received a PI as part of their induction regimen. These transcriptome findings in patient samples are consistent with top hits in our cell line studies, and suggest these transcript changes are driven by Btz resistance. *p*-value by *t*-test. **D.** Immunohistochemistry for CD53 on myeloma plasma cells in bone marrow core biopsies from UCSF patients before and after Btz treatment (*n* = 13) is also consistent with protein-level decrease. H-scoring (see Methods) averaged from two independent hematopathologists (E.R. and S.P.). *p*-value by *t*-test.

Flow cytometry largely validated the extensive surface remodeling uncovered by our proteomic dataset, confirming prominent alterations in CD53, EVI2B, CD50, CD10, and CD151 in both AMO1 (**Fig. 3B**) and RPMI-8226 (**Supp. Fig. 4F)** PI-resistant cells. Our previous analysis indicated that CD53, EVI2B, and CD10 cell surface levels were transcriptionally regulated (**Supp. Fig. 4E**). Therefore, to investigate relevance to myeloma patients, we interrogated mRNA expression in paired pre- and post-first relapse tumor cells in the MMRF CoMMpass dataset (release IA14). 94% of these CoMMpass patients were treated with PI as part of their induction regimen. Consistent with our proteomic findings, in relapsed myeloma patient tumor cells we found significant transcript decreases of *CD53* and *EVI2B,* while *MME* (CD10) showed a significant increase (**Fig. 3C**). *CD151* showed a non-significant increase, though in the context of post-transcriptional regulation protein-level increase could potentially be higher. We further developed an immunohistochemistry (IHC) assay for CD53, confirming decrease on plasma cells in bone marrow core biopsies *(n* = 13) in paired diagnosis and relapse specimens after a PI-containing regimen (**Fig. 3D**). We further noted that these markers appeared to carry prognostic relevance at diagnosis in the CoMMpass cohort (**Supp. Fig. 4G**). From a biological perspective, alterations in these surface proteins may influence the phenotype of PI-refractory myeloma. For example, both CD53 (38) and CD151 (39) may impact tumor interaction with the microenvironment. Taken together, these markers warrant further investigation in the context of assessing and monitoring PI resistance.

### Lenalidomide evolved resistance leads to increased CD33 and CD45/PTPRC on myeloma cells

We further probed surface proteomic changes in the context of evolved resistance to lenalidomide (Len), a thalidomide analog used in both the standard front-line regimen for myeloma patients and maintenance monotherapy. To our knowledge, surface changes resulting from Len resistance have not been previously characterized. We performed CSC proteomics on OPM2 and H929 cell lines *in vitro-evolved* to become resistant to lenalidomide (LenR) relative to their WT counterparts (40). While we found broad surface proteomic changes in both cell lines, there was relatively little overlap between the two (**Fig. 4A**). The most notable common signature was increased CD33 and CD45/PTPRC in both LenR cell lines (**Fig. 4A**). Examining CoMMpass data, we confirmed both *CD33* and *PTPRC* transcripts to be significantly increased at first relapse vs. diagnosis in patient tumor cells (**Fig. 4B**). Plasma cell expression of either of these markers has already been proposed as a poor prognostic factor in newly-diagnosed myeloma (41, 42), potentially consistent with more aggressive disease biology after Len resistance. In terms of therapeutic targeting, while CD45 is expressed at high levels on essentially all non-plasma cell leukocytes, CD33 is a well-known surface target enriched in myeloid malignancies (43). However, by CSC profiling, CD33 shows low mass spectrometric intensity on myeloma plasma cells (**Supp. Dataset 1**), suggestive of low surface protein expression. Furthermore, per Human Blood Atlas data *CD33* mRNA expression on B-lineage cells is expected to be much lower than that on myeloid cells (**Supp. Fig. 4H**). “ On target, off tumor” toxicity on non-malignant cells expressing CD33 is already a considerable concern in treatment of myeloid leukemias (44). Taken together, targeting CD33 to selectively eliminate Len-resistant myeloma is unlikely to be straightforward. However, there may be opportunities to generate dual-targeting cellular therapies or other immunotherapeutics to take advantage of this increased CD33 expression in Len-resistant disease.

**Figure 4.**
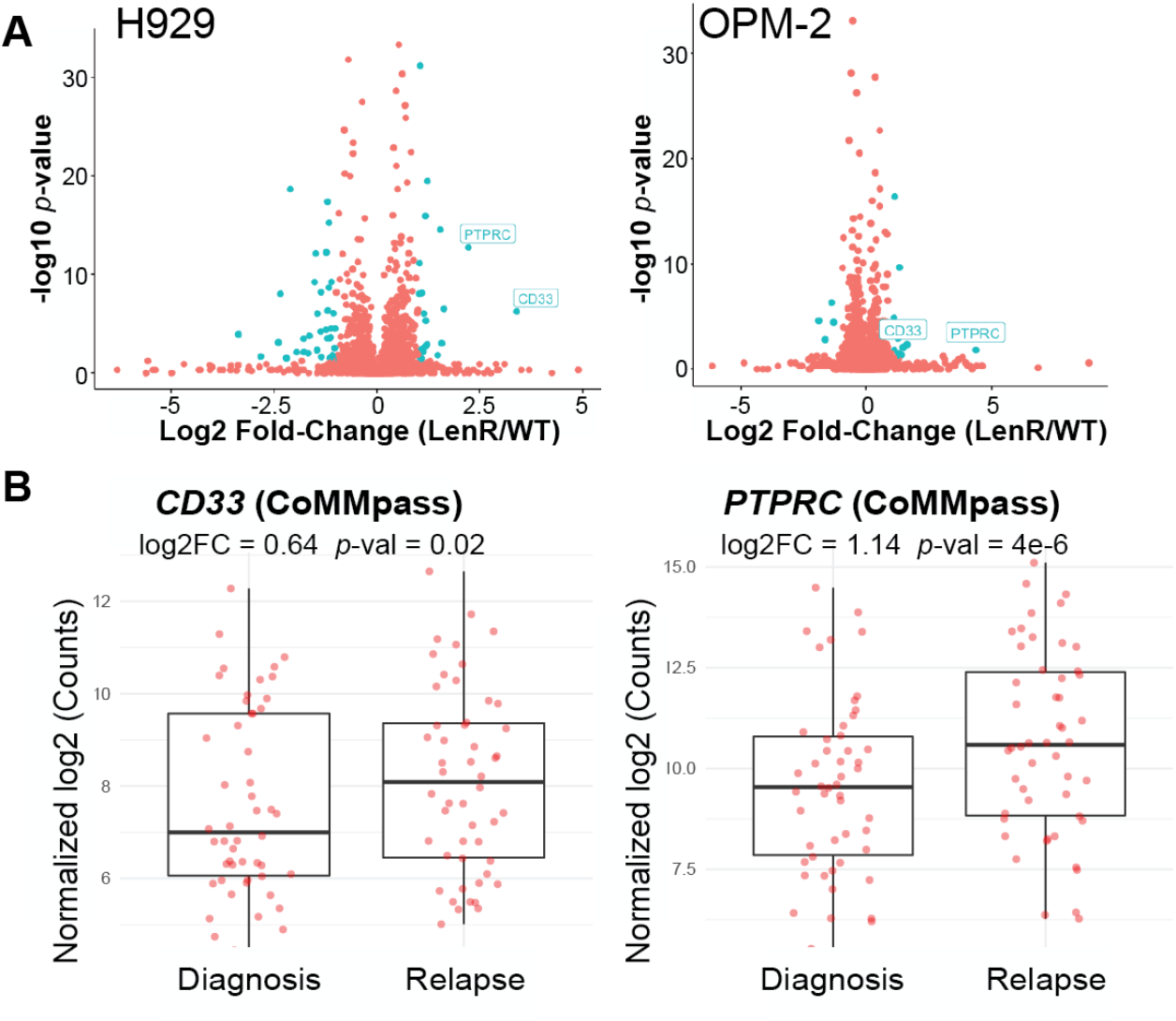
Surface proteomic signatures of lenalidomide resistance. **A.** *In vitro-* evolved lenalidomide-resistant H929 and OPM-2 lines were analyzed by cell surface proteomics with comparison to parental lines by SILAC quantification (*n* = 4; heavy and light channels swapped for two replicates each). Significantly-changed proteins in blue (log2-fold change >|1|;*p* < 0.05), with only CD33 and PTPRC/CD45 showing common changes between the two lines **B.** MMRF CoMMpass patient transcript data confirms significant increase in *CD33* and *PTPRC* at first relapse versus diagnosis, suggesting that increases in these surface proteins is driven by IMiD resistance.

### Short term drug treatment leads to divergent surface profiles from evolved resistance

Another potential strategy to take advantage of the myeloma surfaceome is to consider co-treatment approaches of small molecules and immunotherapies. For example, our group (45) and others (46–48) have used small molecules to increase expression of CD38 on myeloma plasma cells to enhance efficacy of daratumumab. Other examples include small molecule treatment to boost surface BCMA in myeloma (49, 50) or CD22 in B-ALL (51). Furthermore, it has been suggested that acute responses to PI treatment, in particular marked chaperone upregulation and ER stress response, may translate into mechanisms of long-term cellular adaptation and resistance (52–54). We therefore examined the effects of short-term (48 hr) Btz and Len treatment on the myeloma surfaceome (**Supplementary Dataset 5**).

We first noted that surface protein changes after acute Btz treatment at 7.5 nM for 48 hr in RPMI-8226 cells, when compared to BtzR vs. WT data aggregated across cell lines, did not show significant correlation *(R* = −0.056; *p* = 0.058) (**Fig. 5A**). The most apparent commonalities were in down-regulated proteins, including CD53, NCAM2, and SEMA4A. While we did not observe an increase in any current immunotherapy targets after acute Btz treatment, we surprisingly noted a decrease in surface BCMA in RPMI-8226 cells (**Fig. 5B**). We further confirmed this finding in another cell line, MM.1S, by flow cytometry (**Supp. Fig. 5A-B**).

**Figure 5.**
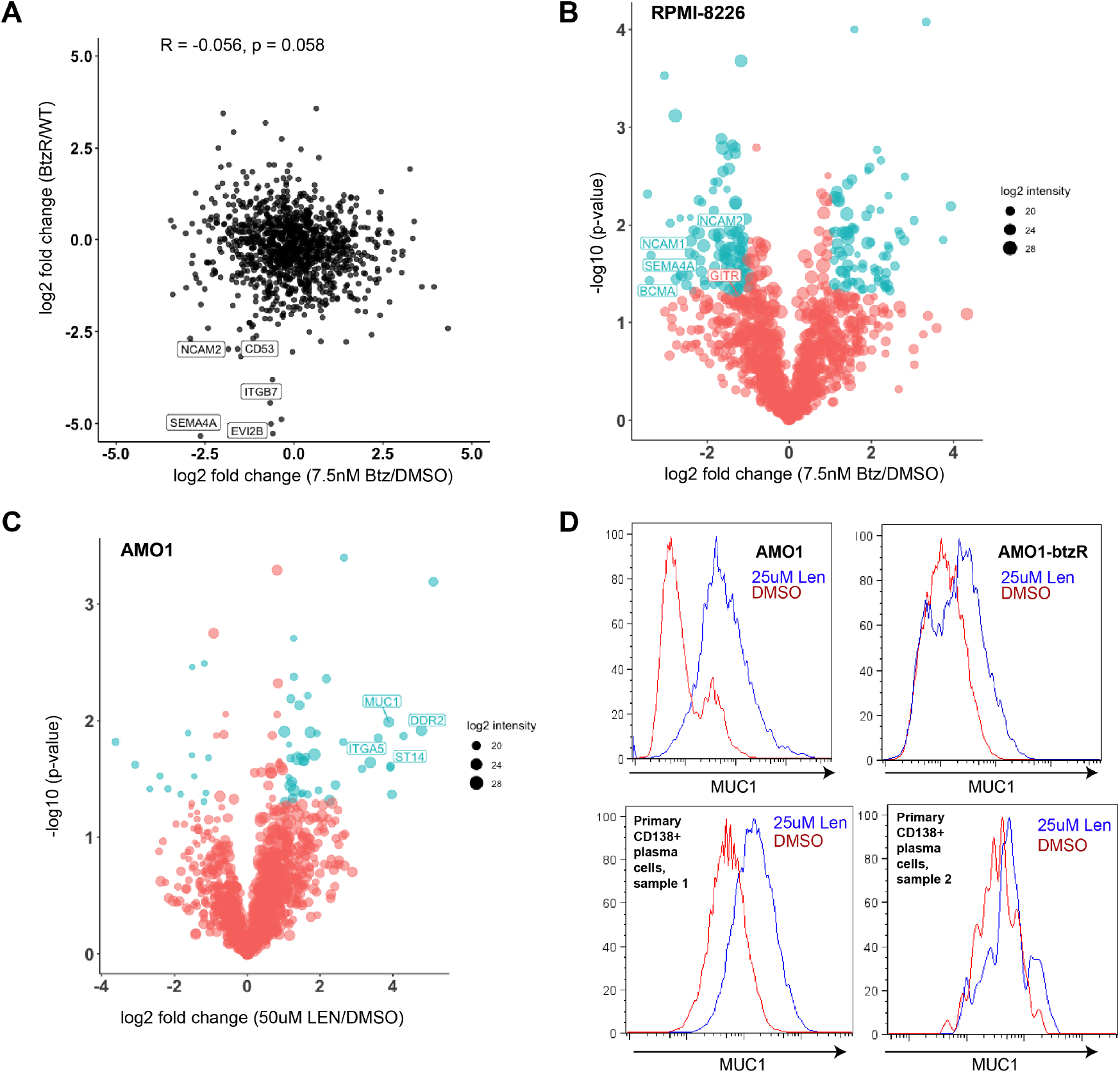
Characterizing myeloma surface proteomic changes in response to acute drug treatment. **A.** Correlation in surface proteomic profile between acute Btz treatment in RPMI-8226 cells (7.5 nM, 48 hrs, *n* = 3) vs. DMSO *(n* = 4) and aggregate BtzR cell line data (as in Fig. 3A) vs. parental. Common changes are observed in some downregulated surface proteins but no significant overall correlation is observed. **B.** Volcano plot of RPMI-8226 cells treated for 48 hr with 7.5 nM bortezomib, highlighting significantly changed proteins (log2-fold change >|1|;*p* < 0.05 in blue). **C.** Similar plot as in B., for 48 hour treatment with 50 μM Lenalidomide in AMO-1 cells. **D.** Validation of increase in surface MUC1 in plasma cells in response to 25 μM Lenalidomide treatment, in both AMO-1, AMO-1 BtzR, and two primary patient bone marrow aspirates (selected on CD138+ plasma cells).

We similarly investigated short term treatment with Len. After 48 hr of 50 uM treatment in AMO-1 we again found broad alterations to the surfaceome (**Fig. 5C**). We did not observe any alteration in CD33 or PTPRC/CD45. However, among the most upregulated proteins was Mucin-1 *(MUC1),* a known therapeutic target in multiple myeloma (55). By flow cytometry we confirmed MUC1 surface changes after short term Len in both AMO-1 as well as primary CD138+ plasma cells from two patients (**Fig. 5D**). Additionally, in RPMI-8226 cells, mass spectrometry and flow cytometry identified GITR, a potential tumor suppressor in myeloma (63), as upregulated under Len treatment (**Supp. Fig. 5C-D**).

We further probed treatment with two other agents targeting protein homeostasis that have been preclinically investigated in myeloma: the p97 inhibitor CB-5083 (56) and the allosteric HSP70 inhibitor JG342 (57, 58) (**Supp. Fig. 5E-G**). We found that surface proteomic signatures were largely unique to both these agents, showing minimal overlap with PI treatment.

Intriguingly, myeloma surface markers show wide variability in expression changes when aggregated across drug treatment conditions (**Supp. Fig. 5H**). For example, TXNDC11 and CCR10 protein levels appear stable across drug treatments in comparison to canonical myeloma markers BCMA and CD138/SDC1. Some proteins, including ITGB6, consistently exhibit large changes in surface expression after drug treatment, suggesting that they might be regulated by common stress pathways in myeloma cells.

Together, these findings confirm that acute drug treatment and long-term resistance lead to rare commonalities but in general largely divergent effects on the myeloma surfaceome. Furthermore, our results suggest that in some settings Btz cotreatment may decrease efficacy of BCMA-targeted therapy, whereas Len co-treatment may enhance utility of MUC1-directed treatments.

### “Micro” method for streamlined surface proteomics

Ideally, our surface proteomic analyses here would be performed on primary myeloma tumor specimens instead of cell lines. However, one significant limitation of the standard CSC method is the requirement for large sample inputs, typically 30e6-200e6 cells (4, 11). In other cancers, this requirement has largely restricted surface proteomic analyses to model systems (20, 59). To our knowledge, no surface proteomic profiling has previously been performed using primary myeloma tumor cells.

We sought to develop a miniaturized CSC method amenable to reduced sample inputs, thereby allowing for more ready application to primary samples. We reasoned that adapting the “InStageTip” strategy could meet this need (60). This “one pot” approach, which minimizes sample losses during processing by eliminating many steps of handling and vessel transfer, was previously shown to allow for large decreases in shotgun proteomic sample input on whole-cell lysates.

In our adaptation of this method for cell surface proteomic profiling, following surface biotinylation of live cells, streptavidin bead capture, and bead washes, all steps are performed in a single P200 tip (**Fig. 6A**). We first applied this “micro” method to RS4;11 B-ALL cells, comparing 1e6 cellular input to 30e6 input in the standard (“macro”) method. With the micro method, by MaxQuant analysis we were able to quantify 662 and 645 proteins in each replicate, compared to 725 and 774, respectively, with the standard macro method with 30x greater input. We further saw consistent quantification between the micro and macro method (*R* = 0.84) and excellent quantitative reproducibility between biological replicates of the micro method on 1e6 cells (*R* = 0.98) (**Fig. 6B-C**).

**Figure 6.**
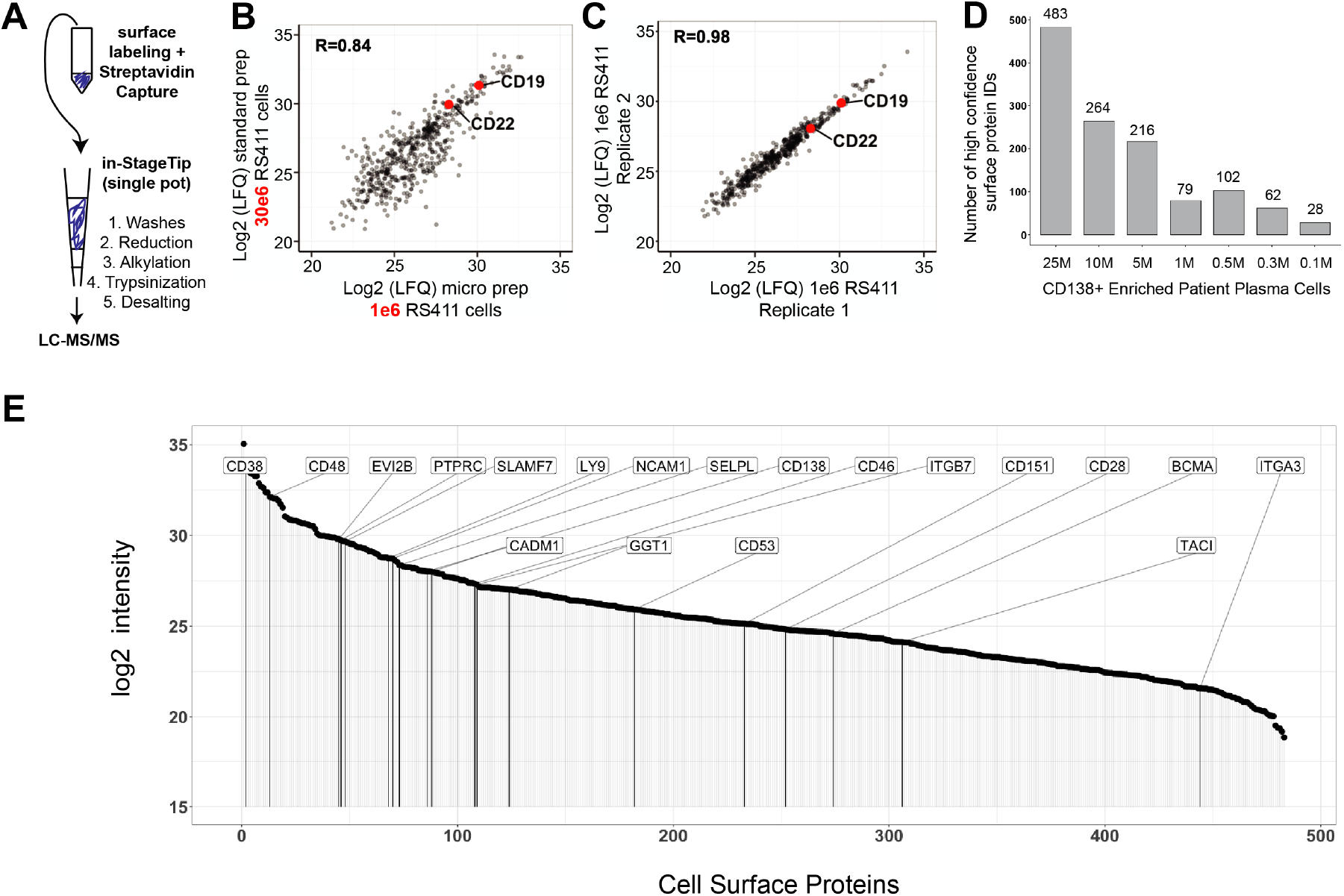
“Micro” protocol for cell surface proteomics. **A.** Schematic of “micro” sample preparation method using an In-StageTip approach for all steps after surface glycoprotein biotinylation on live cells. **B.** Quantitative comparison of LFQ intensity for identified proteins using 30e6 cells in the standard, “macro” protocol, versus 1e6 cells using our “micro” protocol. LFQ values averaged from *n* = 2 biological replicates per preparation method; performed with RS4;11 B-ALL cells. **C.** The micro method demonstrates excellent reproducibility across both biological replicates at 1e6 cell input. **D.** Total number of glycoproteins identified using the “micro” method at various cell inputs, on CD138+ tumor cells isolated from a relapsed/refractory myeloma patient with a malignant pleural effusion. **E.** Cell surface protein intensities in the 25e6 input condition from (D). The large majority of antigens we point out in this work as possibly clinically relevant are also detected in the primary patient samples; several exceptions include CCR10, which as a GPCR likely is not captured efficiently enough to be found by proteomics on this primary sample, CD10, and CD33.

Encouraged by these results, we next evaluated the head-to-head performance of the micro vs. macro method in AMO-1 myeloma cells across a range of inputs (30e6, 10e6, 1e6 cells). In this follow-up experiment, the benefits of the micro prep on protein yield were not as pronounced, with only a modest increase in membrane protein identification with 1e6 cells versus the standard macro prep (**Supp. Fig. 6A**). Furthermore, we noted a marked drop-off in membrane protein identifications from 30e6 to 1e6 cells with micro method, with only 40% and 35% of membrane protein IDs at the lower input in each replicate, respectively (**Supp. Fig. 6A**).

Even though we could not consistently obtain a dramatic increase in membrane protein identifications at the lower sample input, it still appears that the micro method carries at least some advantage in protein identifications over the standard macro protocol on low sample inputs. Furthermore, the “one pot” approach markedly streamlines the overall workflow by eliminating additional steps requiring transfer of the sample from one vessel to another.

We therefore attempted to apply our micro CSC method to primary myeloma plasma cells. We titrated cellular inputs obtained from a relapsed patient with a malignant pleural effusion, from which we isolated ~50e6 CD138+ plasma cells with magnetic bead selection. While we found a marked decrease in membrane protein identifications from 25e6 to 1e6 primary cells (**Fig. 6D**), the micro approach on 5e6 cellular input still enabled quantification of 327 membrane-annotated proteins, of which 216 were high confidence plasma membrane proteins, on this primary tumor sample (**Fig. 6E, Supplementary Dataset 6**). Furthermore, the micro “one pot” approach on 25e6 cellular input quantified 868 total membrane-annotated proteins, of which 483 were high confidence plasma membrane proteins (**Fig. 6E**, **Supplementary Dataset 6**), and identified essentially all of the antigens of interest previously identified in our cell line studies. We further found a strong positive correlation (*R* = 0.51; *p* < 1e4) between surface protein quantification in myeloma cell lines and in this primary sample, with very few surface proteins found on the primary sample not identified in myeloma cell lines (**Supp. Fig. 6B**). This overall similarity to the surface proteomic landscape of a primary tumor underscores the utility of our profiling in myeloma cell lines above.

## DISCUSSION

Here we used a proteomic approach to delineate the cell surface landscape of multiple myeloma plasma cells. Our results characterize the distinct myeloma cell surface repertoire as well as define surfaceome remodeling in the context of resistance to and acute treatment with commonly used small molecule therapeutics. Our findings suggest new immunotherapy targets in this disease, reveal possible biomarkers of resistance, and provide a unique resource for probing plasma cell biology given the numerous physiologic processes governed by cell surface proteins. Furthermore, we modify a common cell surface proteomic method which, with further refinement, may more readily expand “surfaceome” profiling to primary tumor samples.

Notably, our dataset is valuable as it extends to far more proteins simultaneously than are accessible by flow cytometry or CyTOF. On the other hand, for surface proteins present at low copy number, or with no glycosylation, sensitivity of detection by CSC proteomics is certainly lower than these other approaches. Furthermore, we cannot exclude the possibility of false positive hits due to background intracellular protein labeling. Therefore, this broad dataset ultimately serves as an important complement to targeted antibody-based approaches.

We believe that particular utility of this resource comes from its ability to identify surface proteomic signatures that could not be predicted from RNA-seq alone. For example, our identification of surface proteins most highly expressed on plasma cells (**Fig. 2C-E**) cannot be directly extrapolated from transcriptome data. Furthermore, we find evidence of potential post-transcriptional regulation of many surface proteins in the drug-resistant setting (**Supp. Fig. 4E**).

We further integrate our proteomic data with RNA-based datasets to assist in the identification of new immunotherapy targets in myeloma (**Fig. 2A**). Our two hits we specifically examine, CCR10 and TXNDC11, while promising, still require significant additional validation as possible targets. We note that given prior data suggesting ER-localized biology, TXNDC11 in particular may not have been considered as a possible immunotherapeutic target without our proteomic dataset in hand. Other antigens emerging from our scoring system may also have significant promise and stand as a resource for others in the community. In parallel, modulating CAR binding avidity has recently been proposed as a method to precisely tune CAR-T activation (61). Our proposal to use CD48/SLAMF2 binding to increase avidity, and thereby extend functional activity, of BCMA CAR-Ts requires future experimental exploration.

We do note that as we were finalizing this work, another group also performed surface proteomics on a series of myeloma cell lines, as well as targeted whole-cell lysate proteomics from primary myeloma samples (62). In their limited analysis they point out several surface proteins expressed on plasma cells that are being targeted in other malignancies, including CD5, CD147 *(BSG),* LY75 *(LY75),* CD98hc *(SLC3A2),* and CD166 *(ALCAM).* However, based on our scoring metrics (**Supplementary Dataset 2**), all of these appear to be poor cellular therapy targets (score <8) for myeloma due to low plasma cell specificity and high likelihood of on-target, off-tumor toxicity. We believe that our integrated bioinformatic analysis provides a much more robust resource for the development of new myeloma therapies.

In terms of study limitations, due to the sample input required for cell surface proteomics, the large majority of our surface profiling is performed in myeloma cell lines. While we know these models are only partially representative of true patient tumors (63), our identification of nearly all canonical myeloma surface markers and immunotherapy targets on cell lines (**Fig. 1C**; **Fig. 6E**; **Supp. Fig. 6B**) increases confidence in the broader applicability of our findings. In addition, many of the drugresistance or drug-treatment signatures we identified were consistent with those observed in primary patient tumors, either at the transcript (MMRF CoMMpass) or protein (our studies here) level. However, validating the clinical utility of novel surface biomarkers of drug resistance, as well as the potential relevance to small molecule or immunotherapeutic co-treatment strategies (for example, the combination of lenalidomide and MUC1-directed therapy) awaits further investigation.

To this point above, broadly extending surface proteomic profiling to primary plasma cells would certainly be a boon for myeloma research. Our miniaturization of the CSC method provides a small step toward allowing more routine profiling of primary samples, but further optimization is clearly needed. Recently, a method utilizing automated liquid handlers was described, obtaining high-quality data from only 1e6 murine B-cells (64). While this approach is not necessarily optimal as it requires specialized equipment only available in a handful of labs, such approaches may demonstrate a way forward for more routine “surfaceome” profiles of patient plasma cells. However, we still believe that the “micro” method will carry advantages for other groups performing surface proteomics given a streamlined protocol and no apparent disadvantages versus the macro method. Notably, we used our adapted micro method to proteomically profile the cell surface of a myeloma primary sample for the first time. Overall, our data supports the notion that our resource remains highly useful for generating hypotheses and motivating further investigation in myeloma.

In conclusion, we provide a unique resource for the myeloma research community, identifying new surface markers with potential biological, diagnostic, and therapeutic relevance. Through further method advancement, our goal is to ultimately bring this powerful approach to more widespread use in basic, translational, and clinical studies in myeloma.

## Supporting information

Myeloma Leukemia Blymphoblast Mass Spec

Target Scoring

PI Resistant Myeloma Mass Spec

PI Resistant Myeloma RNA

Small Molecule Mass Spec

Myeloma Lenalidomide Resistant Mass Spec

Myeloma Patient Sample Micro Proteomics

Membrane Protein Lists

## ACKNOWLEDGEMENTS

We thank the UCSF myeloma patients and their families who allowed their tissue specimens to be utilized for research in this study. This research was supported by R01 CA226851, DP2 OD022552, the Gabrielle’s Angel Foundation for Cancer Research, and the UCSF Stephen and Nancy Grand Multiple Myeloma Translational Initiative (to A.P.W.), and Celgene UCSF A125812, P41 CA196276, R35 GM122451, and the Harry and Dianna Hind Professorship (to J.A.W.).

## AUTHOR CONTRIBUTIONS

I.D.F., S.T.T, J.A.W., and A.P.W. conceived and designed the study. I.D.F., B.P.E., S.T.T, Y-H.L., M.N., K.K.L., and P.C. performed experiments and analyzed data. M.H. analyzed data. E.R., S.V., and S.P. performed and analyzed immunohistochemistry. A.L.-G., L.B., and C.D. contributed reagents. N.S., S.W.W., J.L.W., and T.G.M. enrolled patients in tissue banking protocol and provided primary patient samples. I.D.F. and A.P.W. wrote the manuscript with input from all authors.

## CONFLICT OF INTEREST

A.P.W. is a member of the Scientific Advisory Board and holds equity stakes in Indapta Therapeutics and Protocol Intelligence, LLC. J.A.W. is on the Scientific Advisory Board and holds equity stakes in the following companies with oncology interests: Soteria Biotherapeutics, Jnana Therapeutics, Inception Therapeutics, and Inzen Therapeutics, and holds a sponsored research agreement with Bristol Myers Squibb. S.W.W. has received research funding from Janssen, GlaxoSmithKline, Bristol Myers Squibb, Genentech, and Fortis, and served as a consultant to Amgen. T.G.M. has received research funding from Sanofi, Janssen, and Amgen and has served as a consultant to GlaxoSmithKline. A. L.-G. is an employee and shareholder of Celgene/Bristol Myers Squibb. P.C. is currently an employee and shareholder of Roche/Genentech but was solely employed by UCSF during her participation on this project. The other authors declare no relevant conflicts of interest.

## METHODS

### Cell Lines

All cell lines were grown in RPMI-1640 media with 10% FBS. PI-resistant cells derived from cell lines AMO-1, L363, RPMI-8226, and ARH-77 were grown in 90 nM Bortezomib (Btz) or Carfilzomib (Cfz) as previously described (65). Lenalidomide-resistant cell lines H929 and OPM-2 were grown in increasing concentrations of lenalidomide and removed from drug for 5-7 days, as previously described (40), before use in cell surface proteomics. Cell lines were verified by DNA short tandem repeat testing and assessed as mycoplasma negative.

### Myeloma Patient Sample Processing

Patient bone marrow samples (BM) were processed as previously described (66). Fresh de-identified primary myeloma patient BM samples were obtained from the UCSF Hematologic Malignancies Tissue Bank in accordance with the UCSF Committee on Human Research-approved protocols and the Declaration of Helsinki. BM mononuclear cells were isolated by density gradient centrifugation with Histopaque-1077 (Sigma Aldrich), and washed with 10 mL DPBS 3 times. For small molecule perturbation experiments, mononuclear cells were plated in IL-6 supplemented medium (RPMI1640, 10% FBS, 1% penicillin/streptomycin, 2 mM glutamine, with 50ng/mL recombinant human Il-6 (ProSpec)) at 2e5 cells per well in a 96-well plate and incubated at 37°C, 5% CO2 overnight prior to drug treatment and processing for flow cytometry as described below. For mass spectrometry, isolated mononuclear cells were taken directly from histopaque isolation into CD138 magnetic bead isolation and surface glycoprotein labeling as described below.

### Flow Cytometry

Cells were resuspended in FACS buffer (5% FBS in D-PBS) and stained with antibodies for 1-2 hours on ice, washed with FACS buffer, and then resuspended in FACS buffer. For experiments where live cells populations were studied, cells were resuspended in FACS buffer. For analyses including live/dead stains, cells were resuspended in FACS buffer containing Sytox Green or Sytox Red (Thermo). For drug treatment experiments, cells were plated into 96 well plates and treated for 48 hours with compounds unless otherwise noted. For quantitative flow cytometry, antibodies used were FITC Mouse antihuman CD48 (BD Biosciences, clone TU145), FITC Mouse anti-human CD38 (BD Biosciences, clone HIT2), FITC Mouse IgG1k isotype control (BD Biosciences, clone 27-35), and unstained respectively. using calibrated beads (Bangs Labortories), 200 μL of FACS buffer + 1 drop of beads per well (one wells for “blank” beads, and other for FITC-Beads). Calibration curve for quantification beads were performed on the Quantitative software QuickCal v2.3. (Bangs Laboratories, Inc.) before FITC analysis on multiple myeloma cell lines. The following antibodies were used: CD138 (BD Biosciences, 562097, 552026), BCMA (BD Biosciences, 552026, BioLegend 357504), CD53 (BD Biosciences, 555508), CD10 (BD Biosciences, 561002), CD151 (BD Biosciences, 556057), CD50/ICAM3 (BD Biosciences, 555958), ITGB7 (BD Biosciences, 555945), GITR/CD357 (BioLegend 311604), MUC1/CD227 (BD Biosciences, 559774), CCR10 (BD Biosciences, clone 1B5). Isotype antibodies for markers were ordered from BD Biosciences and BioLegend as per manufacturer recommendations.

### Drug treatments

For cell surface proteomics experiments, 25e6 cells were seeded at 1e6/mL in T75 flasks and treated with compounds for 48 hours at the LD25 dose. RPMI-8226 cells were treated with 50uM Lenalidomide (Sigma CDS022536), 7.5nM Bortezomib (SelleckChem), or 300nM CB-5083 (gift of Cleave Biosciences). AMO1 cells were treated with 50uM Lenalidomide, 750nM JG-342 (gift of Jason Gestwicki, UCSF), or 250nM CB-5083. For flow cytometry experiments, cells were plated and treated in 96-well plates with the doses described in figure legends for 48 hours prior to flow cytometry analysis unless otherwise indicated.

### Cell surface proteomics sample preparation

For all analyses except lenalidomide-resistant cell lines, 30e6 cells were collected for cell surface proteomics and washed twice with cold PBS. To oxidize glycoproteins, cells were incubated with 1.6mM Sodium metaperiodate (NaIO_4_) for 20 minutes with end-over-end rotation at 4°C in the dark, followed by three washes with ice cold PBS. To label oxidized glycoproteins, cells were incubated with 10mM Aniline and 1mM Biocytin hydrazide (Biotium, 90060) for 1.5 hours at 4°C with end-over-end rotation. Cells were washed three times with ice cold PBS, snap frozen in liquid nitrogen, and stored at −80C prior to processing. Pellets were lysed in 2x RIPA Buffer with HALT Protease Inhibitors (Thermo) and 2mM EDTA and sonicated on ice with occasional vortexing. Lysates were mixed with 500uL Neutravidin (Pierce, 29200) bead slurry and incubated on end over end rotary for 120 minutes at 4°C. Slurry was washed on columns with 50mLs each of three wash buffers as follows: first, 1x RIPA with 1mM EDTA, second, PBS with 1M NaCl, and third, 50mM ammonium bicarbonate with 2M Urea. Bead slurry was transferred to an eppendorf tube with 50mM Tris pH 8.5, 10mM TCEP, 20mM IAA, 1.6M Urea. 2ug Trypsin was added to each sample for overnight digestion at RT. Supernatant contained digested peptides was removed from bead slurry and acidified (pH ~2) prior to desalting on SOLA columns or homemade C18 Stagetips. Eluted peptides were dried down by speedvac and resuspended in 2% ACN, 0.1% FA for mass spectrometry analysis. For lenalidomide-resistant cell line analysis using SILAC, all cell lines were grown were cultured in RPMI SILAC media (Thermo) supplemented with Dialyzed FBS for SILAC (Thermo) containing L-[^13^C_6_,^15^N_2_]lysine and L-[^13^C_6_,^15^N_4_] arginine (heavy label; Thermo) or L-[^12^C_6_,^14^N_2_]lysine and L-[^12^C_6_,^14^N_4_]arginine (light label, Thermo) for 5 passages to ensure full incorporation of the isotope labeling on cells. Briefly, 40×10^6^ Len sensitive and resistant cells were harvested at 80% confluence, mixed at 1:1 cell count ratio, and subjected to the CSC protocol as described above to include all tryptic fragments. All experiments were performed in duplicate in both forward and reverse SILAC labeling scheme such that a total of four biological replicates were analyzed together.

### Macro-vs-Micro Cell Surface Proteomics

AMO1 cells were counted and titrated into 3e7, 1e7, and 1e6 cells for cell-surface labeling. For harvesting, cells were washed twice with cold D-PBS (UCSF CCFAL001). For cell-surface labeling, the samples were re-suspended in 990 μL cold D-PBS and transferred to a 1.5-mL amber tube. Next, they were oxidized by the addition of 10 μL 160mM NaIO_4_ (Thermo 1379822) and incubated on a rotisserie at 4°C for 20 minutes. Three spin washes were performed each with 1 mL cold D-PBS at *300g* for 5 minutes to remove the oxidizing reagent. For chemical labeling, cell pellets were re-suspended in 1 mL cold D-PBS followed by the addition of 1 μL of aniline (Sigma-Aldrich 242284) and 10 μL biocytin hydrazide (Biotium 90060). Samples were incubated at 4°C for 60 minutes on a rotisserie followed by three more spin washes with cold D-PBS. After the final wash, supernatant was removed, and cell pellet were snap frozen and stored in −80°C. Labeling was repeated for duplicate samples for the two protocols.

For both micro and macro protocols, labeled cell pellets were lysed in 500 μL buffer containing 2X RIPA (Millipore 20-188), 1X HALT protease inhibitor (Thermo 78430), and 2 mM EDTA. Lysates were sonicated at 1-Hz pulses for 30 seconds with a probe sonicator and incubated on ice for 10 minutes. To remove precipitates, samples were spun at 17,200*g* for 10 minutes at 4°C. To prepare for enrichment of biotinylated proteins, 100, 80, 60 of High Capacity NeutrAvidin slurry (Thermo PI29204) were washed three times with 1 mL 2X RIPA/1mM EDTA buffer in 2-mL chromatography columns (Bio-Rad 7326008) attached to a vacuum manifold (Promega A7231). Clarified lysates were then transferred to the chromatography column (3e7, 1e7, 1e6 cell inputs into 100, 80, 60 slurry, respectively) and incubated at 4°C for 60 minutes on a rotisserie. For the macro protocol, protein-bound beads were transferred to a 10-mL chromatography column (Bio-Rad 7311550) and washed with the following buffers: 50 mL 1x RIPA/1mM EDTA, 50 mL 1X PBS/1M NaCl, and 50 mL 2M urea/50mM ammonium bicarbonate (ABC). For the micro protocol, beads were transferred to a 2-mL chromatography column and washed with 5 mL of each buffer. For the macro protocol, rinsed beads were dried fully and transferred to a 1.5-mL tube using 100 μL of digestion buffer containing 1.5M urea (VWR 97063-798), 50mM ABC, 10 mM 2-chloroacetamide (VWR 97064-926), and 5mM TCEP (GoldBio TCEP10). For the micro protocol, P200-StageTips were packed with four C18 disks (3M 14-386-2) and activated with 60 μL methanol, 60 μL 80% acetonitrile (ACN)/0.1% formic acid (FA), and twice with 60 μL 0.1% trifluoroacetic acid (TFA) prior to transferring the beads to the tip using 100 μL of the digestion buffer. For protein digestion, trypsin/LysC (Promega PRV5073) was reconstituted to 1 μg/μL with ultrapure water, and added to each sample (1500, 500, 50 ng trypsin for 3e7, 1e7, 1e6 cell inputs, respectively). The macro samples were mixed at 1000 RPM on an orbital shaker while the micro samples were wrapped in parafilm and placed on an end-to-end rotisserie at RT for the overnight digestion.

For the macro protocol, the soluble fraction containing the tryptic peptides was separated from the beads by spinning at 1000*g* for one minute. Samples were then transferred to fresh tube and acidified with 10% TFA to reach a final concentration of 1% TFA. For desalting, C18 SOLA columns (Thermo 03150391) were activated with 500 μL ACN and equilibrated twice with 500 μL 0.1% TFA on a vacuum manifold. Acidified samples were passed through the column twice followed by two washes with 1000 μL 0.1% TFA and one wash with 500 μL 2% ACN/0.1% FA. Finally, peptides were eluted once with 150 μL 80% ACN/0.1% FA and again with 200 μL 80% ACN/0.1% FA. Samples were fully dried by SpeedVac and stored in −80°C.

For the micro protocol, 10% TFA was added to each sample to reach a final concentration of 1% TFA. An initial peptide binding step was performed by spinning the acidified samples at 1000*g* prior to desalting three times with 100 μL 0.1% TFA. Peptides were then eluted twice with 50 μL 80% ACN/0.1% FA. Samples were fully dried by SpeedVac and stored in −80°C.

### LC-MS/MS operation

For all analyses except lenalidomide-resistant cell lines, 1 ug of peptides were injected into a Dionex Ultimate 3000 NanoRSLC instrument with 15-cm Acclaim PEPMAP C18 (Thermo) reverse phase column coupled to a Thermo Q Exactive Plus mass spectrometer. A linear gradient from 2.4% Acetonitrile to 40% Acetonitrile in 0.1% Formic Acid over 195 minutes at flow rate of 0.2uL/min was employed, with an increase to 80% Acetonitrile for column wash prior to re-equilibration. For MS1 acquisition, spectra were collected in data dependent top 15 method with full MS scan resolution of 70,000, AGC target was set to 3e6, and maximum IT set to 100ms. For MS2 acquisition, resolution was set to 17,500, AGC set to 5e4, and maximum IT to 180ms. For lenalidomide-resistant cell line analysis using SILAC, data were collected on the Q-Exactive Plus in data-dependent mode using a top 20 method with dynamic exclusion of 35 secs and a charge exclusion setting that only sample peptides with a charge of 2, 3, or 4. Survey scans were collected as profile data with a resolution of 140,000 (at 200 m/z), AGC target of 3E6, maximum injection time of 120 ms, and scan range of 400 – 1800 m/z. MS2 scans were collected as centroid data with a resolution of 17,500 (at 200 m/z), AGC target of 5E4, maximum injection time of 60 ms with normalized collision energy at 27, and an isolation window of 1.5 m/z with an isolation offset of 0.5 m/z.

### Proteomic data analysis and quantification

For all analyses except lenalidomide-resistant cell lines, mass spectrometry data was processed in Maxquant (12) (version 1.6.2.1) with settings as follows: enzyme specificity as trypsin with up to two missed cleavages, PSM/Protein FDR 0.01, cysteine carbidomethylation as fixed modification, methionine oxidation and N-terminal acetylation as variable modifications, minimum peptide length = 7, matching time window 0.7 min, alignment time 20 min, with match between runs, along with other default settings. Data was searched against the Uniprot Swiss-Prot human proteome (downloaded Sept. 3, 2018). Proteingroups files were exported from Maxquant, filtered to remove contaminants, and filtered for proteins with at least two unique peptides for analysis. Data analysis was performed in Perseus (67) and R (69). Identified proteins were filtered against curated lists of Uniprot-annotated membrane proteins or those identified in the analysis of Bausch-Fluck *et al.* (10) (Supplementary Table 8).

For lenalidomide-resistant cell line analysis using SILAC, peptide search for each individual dataset was performed using ProteinProspector (v5.13.2) against 20203 human proteins (Swiss-prot database, obtained March 5, 2015). Enzyme specificity was set to trypsin with up to two missed cleavage; cysteine carbamidomethyl was set as a fixed modification; methionine oxidation, lysine and arginine SILAC labels were set as variable modifications; peptide mass tolerance was 6 ppm; fragment ion mass tolerance was 0.4 Da; peptide identification was filtered by peptide score of 0.0005 in Protein Prospector, resulting in a false discovery rate (FDR) of <1% calculated by number of decoy peptides included in the database.

Quantitative data analysis was performed using Skyline (68) software with the MS1 filtering function. Specifically, spectral libraries from forward and reverse SILAC experiments were analyzed together such that MS1 features without an explicit peptide ID would be quantified based on aligned peptide retention time. The first four isotopic peaks of precursor ions were then quantified at 50% FWHM defined by ms1 scanning resolution of 140,000 (at 200 m/z). The boundary of peptide elution time was determined by default algorithm and the total peak area was used as the peptide quantification value. An isotope dot product of at least 0.8 (as calculated by Skyline) was used to filter out low quality peptide quantification, and a custom report was generated for further processing and analysis using R. To ensure stringent quantification of the surface proteome, several filters were applied to eliminate low confidence protein identifications. In the tryptic fraction, only peptides with five or more well quantified peptides were included. Each replicate between individual fractions or reverse SILAC labeling agreed with each other, and forward and reverse SILAC datasets were then combined and reported as median log2 enrichment values. All p values reported is Wilcox ranked test for median log2 enrichment SILAC ratio. All data analyses were carried out using R.

### Cell Surface Proteomics on Primary Myeloma Cells

CD138^+^ myeloma cells were isolated from patient peripheral blood using a magnetic bead-based approach (EasySep Human CD138 Positive Selection Kit, StemCell Technologies 17877). The CD138^+^-enriched sample was titrated into 2.5e7, 1e7, 5e6, 1e6, 5e5, 3e5, 1e5 cells for cell-surface labeling and processed to obtain surface peptides using the micro protocol described above.

### CD53 Immunohistochemistry

Decalcified bone marrow core biopsies were obtained from myeloma patients under a UCSF Committee on Human Research-approved protocol. The electronic medical record was evaluated to identify sequential core biopsies performed on the same patient at both diagnosis and at first relapse. Interpretation was performed by two hematopathologists (S.P. and E.R.) blinded to patient identification and relapse status. Final H-score was averaged between pathologists. CD53 staining was assessed on cells with morphology consistent with plasma cells and CD138+ staining on adjacent tissue sections.

For immunohistochemical staining, tissues were fixed in neutral buffered 10% formalin for 24 to 48h, dehydrated with graded alcohols and infiltrated with paraffin wax at 58 degrees in an automated tissue processor. Infiltrated tissue was embedded at 60C to produce FFPE blocks. FFPE blocks were sectioned at 4 microns. Immunohistochemical detection on the unstained FFPE tissue sections was performed on Ventana Medical Systems Discovery Ultra Biomarker Automated Slide Preparation System using alkaline epitope conditioning (Ventana/Roche CC1) at 97C for 32 minutes. Recombinant rabbit monoclonal (clone EPR4342(2) directed to CD53 supplied by Abcam (ab134094) used at 1:200 for 32 minutes at 36C. OmniMap anti-Rb HRP (Ventana 760-4311) was used for chromogenic DAB detection (32 minutes).

### Bioinformatic Analysis of “Locking on” and standalone immunotherapy targets

For “locking on” analysis, LFQ intensity averages, quartiles, and standard deviation were calculated based on information in Supplementary Dataset 1. The 14 potential locking-on targets were compared to transcript levels in MMRF_CoMMpass_IA16a_E74GTF_cufflinks_Gene_FPKM data set.

For standalone immunotherapy targets, the scoring rubric used is described in **Supplementary Table 1**. For COMPARTMENTS predictions of subcellular localization, the data set used was “human_compartment_integrated_full.tsv”, available on https://compartments.jensenlab.org/Downloads and accessed on August 27, 2020. The CCLE database used for this analysis was: “CCLE_RNAseq_rsem_genes_tpm_20180929.txt” and accessed on June 25, 2020. Available on https://portals.broadinstitute.org/ccle/data. For the points assignment, a TPM average values was taken and a non-heme cancer cell line average was calculated. Also, the values from each disease were transformed into log2 values (and the nonheme averages). The nonheme diseases were: Bile Duct, Breast, Chondrosarcoma Colorectal, Endometrium, Esophagus, Ewings Sarcoma, Giant cell tumor, Glioma, Kidney, Liver, Lung NSC, Lung small cell, Medulloblastoma, Melanoma, Mesotelioma, Neuroblastoma, Osteosarcoma, Other, Ovary, Pancreas, Prostate, Soft Tissue, Stomach, Thyroid, Upper Aerodigestive, Urinary Tract. The genes where Multiple myeloma were highest expressed compared to the other diseases were selected and compared to the average value in non-heme tissues. For Human Blood Atlas analysis, the database used was “rna_blood_cell_monaco.tsv” available in the Human Protein Atlas portal https://www.proteinatlas.org/about/download. Accessed on August 27, 2020.

For GTEx analysis, the dataset used was “GTEx_Analysis_2017-06-05_v8_RNASeQCv1.1.9_gene_median_tpm.gct”, accesed on July 16, 2020 and available on https://gtexportal.org/home/datasets. The ESNG gen names were transformed into HUGO nomenclature using gprofiler2 package in RStudio (https://CRAN.R-project.org/package=gprofiler2). We calculated a non-heme tissue average and a heme tissue average. The non-heme tissues were: Adipose Subcutaneous, Adipose Visceral Omentum, Adrenal Gland, Artery Aorta, Artery Coronary, Artery Tibial, Bladder, Brain Amygdala, Brain Caudate basal ganglia, Brain Cerebellar Hemisphere, Brain Cerebellum, Brain Cortex, Brain Frontal Cortex BA9, Brain Hippocampus, Brain Hypothalamus, Brain Nucleus accumbens basal ganglia, Brain Putamen basal ganglia, Brain Spinal cord cervical c1, Brain Substantia nigra, Breast Mammary Tissue, Cells Cultured fibroblasts, Cervix Ectocervix, Cervix Endocervix, Colon Sigmoid, Colon Transverse, Esophagus Gastroesophageal Junction, Esophagus Mucosa, Esophagus Muscularis, Fallopian Tube, Heart Atrial Appendage, Heart Left Ventricle, Kidney Cortex, Kidney Medulla, Liver, Lung, Minor Salivary Gland, Muscle Skeletal, Nerve Tibial, Ovary, Pancreas, Pituitary, Prostate, Skin Not Sun Exposed Suprapubic, Skin Sun Exposed Lower leg, Small Intestine Terminal Ileum, Skin Sun Exposed Lower leg, Small Intestine Terminal Ileum, Stomach, Testis, Thyroid, Uterus, and Vagina. The heme tissues included were Cells-EBV transformed lymphocytes, Spleen, and Whole Blood.

In the end, we merged these 5 data sets, matching by Gene Name, using the controls mentioned above (used across all the process to verify reliability), for 33654 proteins scored in total. The final data set were created with all the points assigned across all the data set, creating a total score which is the sum of the prior points. Then, the proteins were ranked by the higher score in a descending manner (see **Supplementary Dataset 2**).

### Data Availability

Proteomic data generated in this study was deposited to ProteomeXchange via the PRIDE database (69, 70) (PXD022482, PXD022553). Leukemia proteomic data was downloaded from PRIDE (PXD016800). RNA sequencing data generated in this study was uploaded to GEO (GSE160572). Processed leukemia RNA sequencing data was downloaded from GEO (GSE142447).

## SUPPLEMENTARY FIGURES and LEGENDS

**Supplementary Figure 1.**
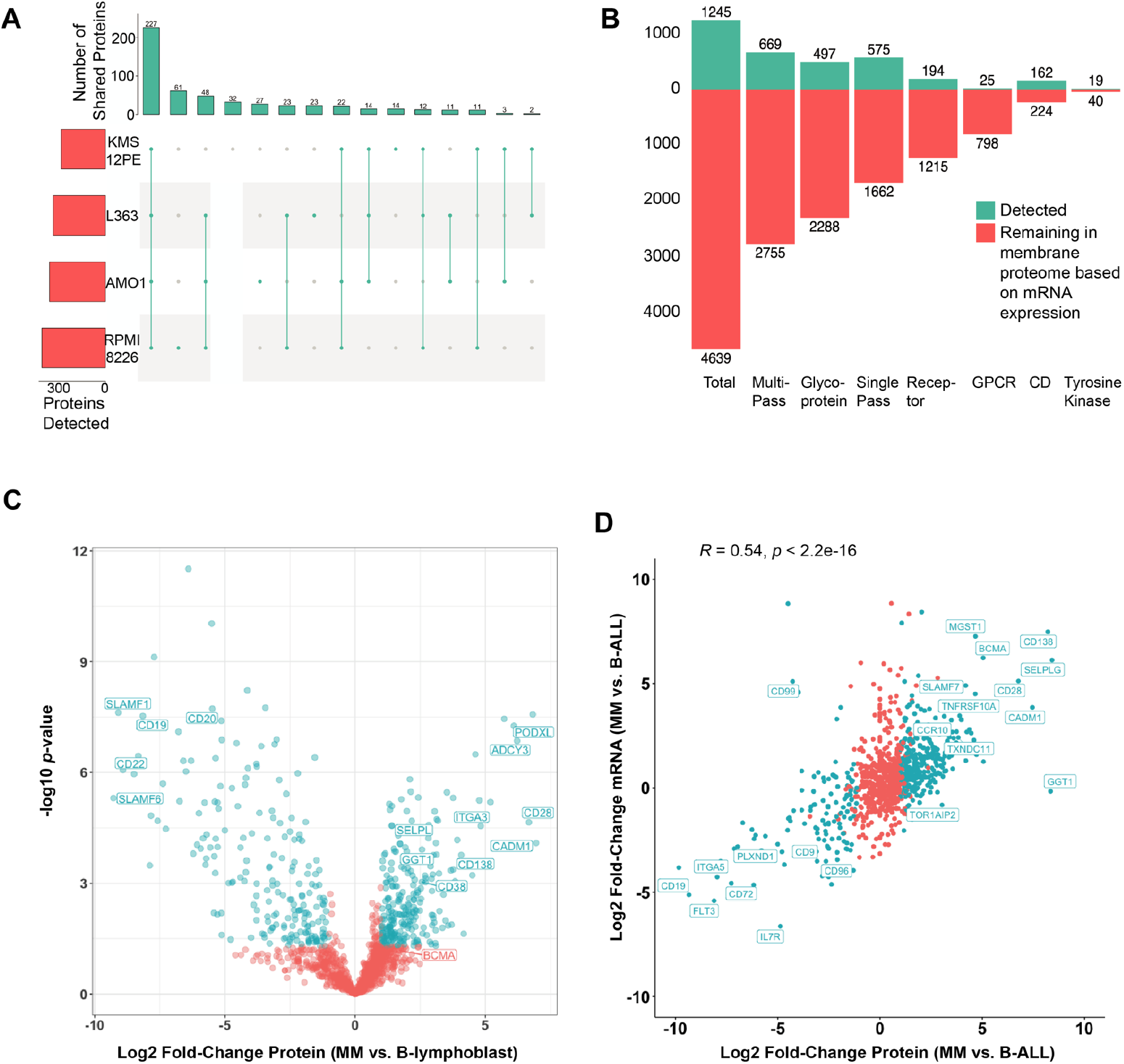
Additional characterization of the baseline myeloma surfaceome. **A.** Quantified captured proteins (minimum 2 peptides per protein) filtered based on highest-confidence plasma membrane localization, from ref. (10). **B.** All genes expressed at mRNA level TPM > 1 in the four analyzed cell lines included (per data available at keatslab.org) were filtered by Uniprot annotation for different protein classes. We then evaluated which fraction were detected by our cell surface capture experiments. For example, in this data we identify 45% of Cluster of Differentiation (CD) markers predicted to be present on myeloma cells based on mRNA expression, but only 6% of GPCRs. **C.** Analogous to Fig. 1E, we compared our myeloma surface profiles across myeloma lines (AMO1, L363, RPMI-8226, and KMS12PE) to the two B-lymphoblastoid lines examined (ARH-77 and EBV-immortalized normal donor-derived) to identify markers that most-distinguish plasma cells. Significantly-changed proteins noted in blue (log2-fold change >|1|;*p* < 0.05). **D.** RNA-seq data on B-ALL and myeloma cell lines (as in Fig. 1E) was also analyzed for log2-fold changes between cell types and compared to log2-fold changes from proteomics. This analysis identifies proteins that may be under post-transcriptional regulation of surface expression, particularly with significant increases in mRNA but no detected change in captured protein. Proteins labeled in blue identical to those found to be significant in volcano plot of Fig. 1E. Pearson correlation and associated *p*-value of significance shown.

**Supplementary Figure 2.**
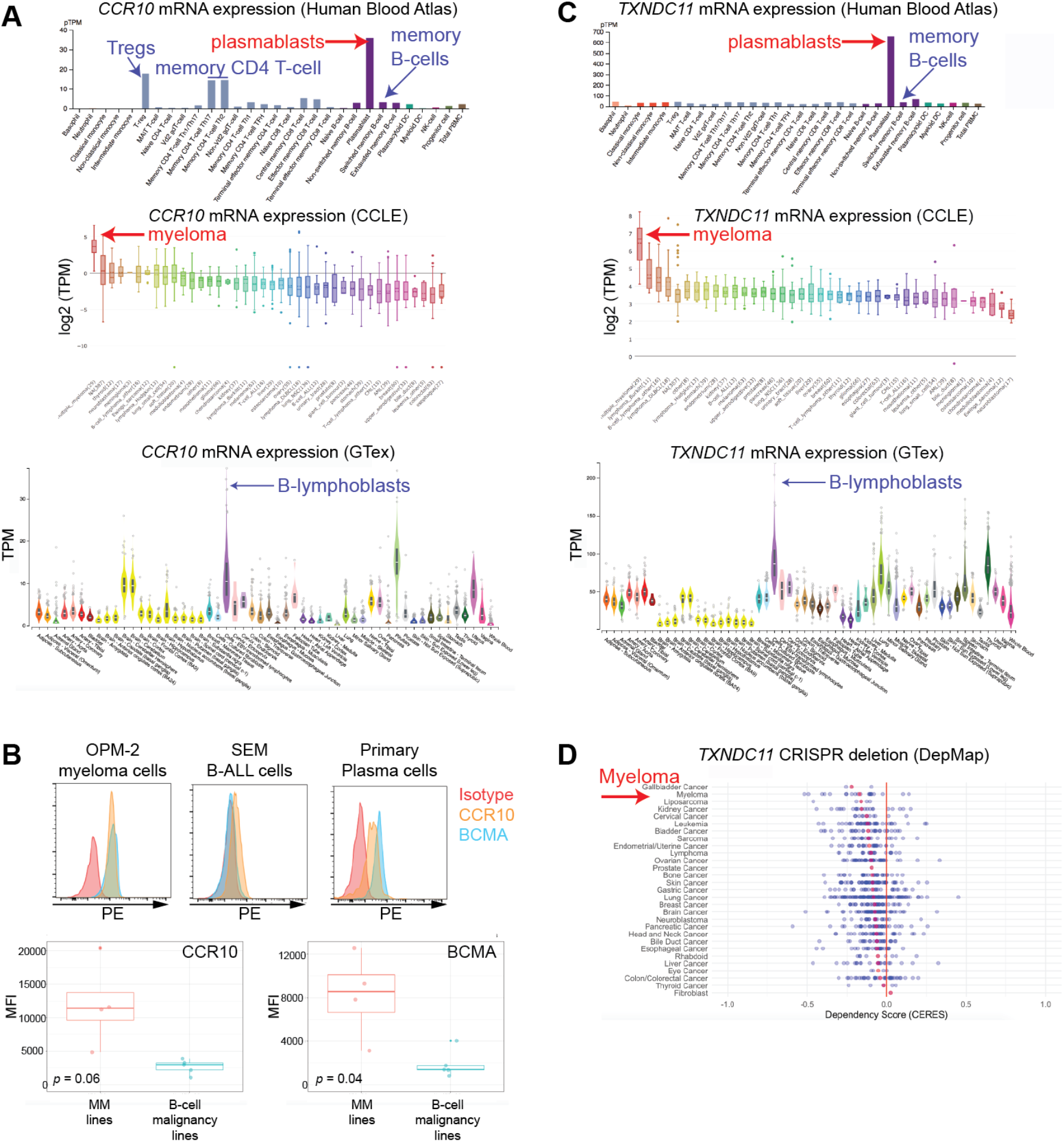
Additional characterization of CCR10 and TXNDC11 as potential myeloma immunotherapy targets. **A.** mRNA transcript data from the Human Blood Atlas (top) demonstrates that among hematopoietic cells *CCR10* is most highly expressed in plasmablasts (we note that long-lived plasma cells are unfortunately not included in this dataset; plasmablasts used as closest proxy). Cancer Cell Line Encyclopedia (CCLE) (middle) data supports that on average myeloma cell lines express >20x more CCR10 mRNA than any other tumor cell type. GTex (bottom) data does suggest some low level CCR10 expression in non-hematopoietic tissues; B-lymphoblast expression can be compared to memory B-cell expression in the Human Blood Atlas. Plasmablast expression is therefore expected to be at least 20x higher than other non-hematopoietic tissues, supportive of a therapeutic index. **B.** Flow cytometry analysis confirms significantly higher CCR10 expression on myeloma plasma cells (OPM-2, MM.1S, ANBL-6, AMO-1) than B-cell malignancy cell lines (SEM, RS411 (B-ALL), HBL-1, OCI-LY10, TOLEDO (B-cell lymphoma)), with similar increase as seen for BCMA. Analysis of a primary patient plasma cell (CD19-/CD138+/CD38+) specimen confirms CCR10 expression *(top right).* **C.** Similar analysis as in A., illustrating highly enriched *TXNDC11* expression on plasmablasts, highly increased expression in myeloma plasma cells versus any other cancer cell type in the CCLE, and moderate expression in other non-hematopoietic tissues. **D.** Data from the Cancer Dependency Map (depmap.org; Avana Public 20Q3) indicate that myeloma cell lines are among the most genetically dependent on CRISPR deletion of this gene, as noted by lowest CERES score when averaged across all included tumor cell lines.

**Supplementary Figure 3.**
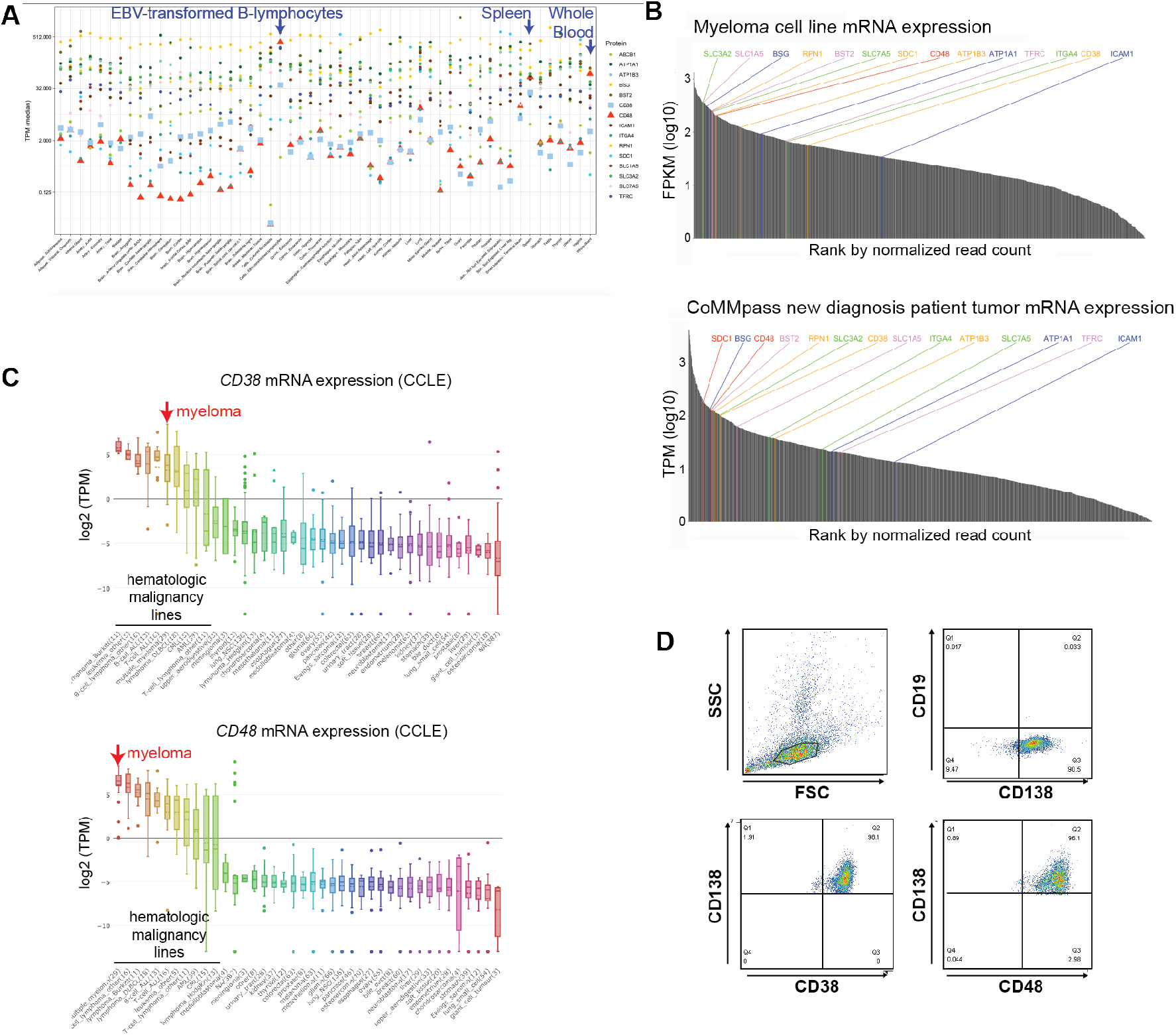
Additional characterization of high abundance surface antigens for a potential “locking on” strategy. **A.** Plot of “locking on” candidate (as in Fig. 2D) transcript expression in GTex suggests that most genes have significant expression in non-hematopoietic tissues, with the exception of CD38 and CD48. **B.** mRNA expression of all genes corresponding to proteins identified in our surface proteomic datasets were ranked according to average abundance (measured in TPM) in both cell lines (keatslab.org dataset) and primary CD138+ tumor cells (MMRF CoMMpass release IA15). Potential “locking on” candidates are highlighted. **C.** CCLE data highlights that myeloma cell lines express the highest average *CD48* mRNA, whereas several other hematopoietic malignancy cell types express greater *CD38* than myeloma cells. These data also confirm that non-hematopoietic cells do not appear to express these antigens. **D.** Illustration of flow cytometry gating strategy to identify primary patient tumor cells for use in absolute quantification of CD38 and CD48. Mononuclear cells in patient bone marrow aspirate were gated on singlet, live lymphocytes in the SSC/FSC plot and then characterized as CD138+/CD19-plasma cells.

**Supplementary Figure 4.**
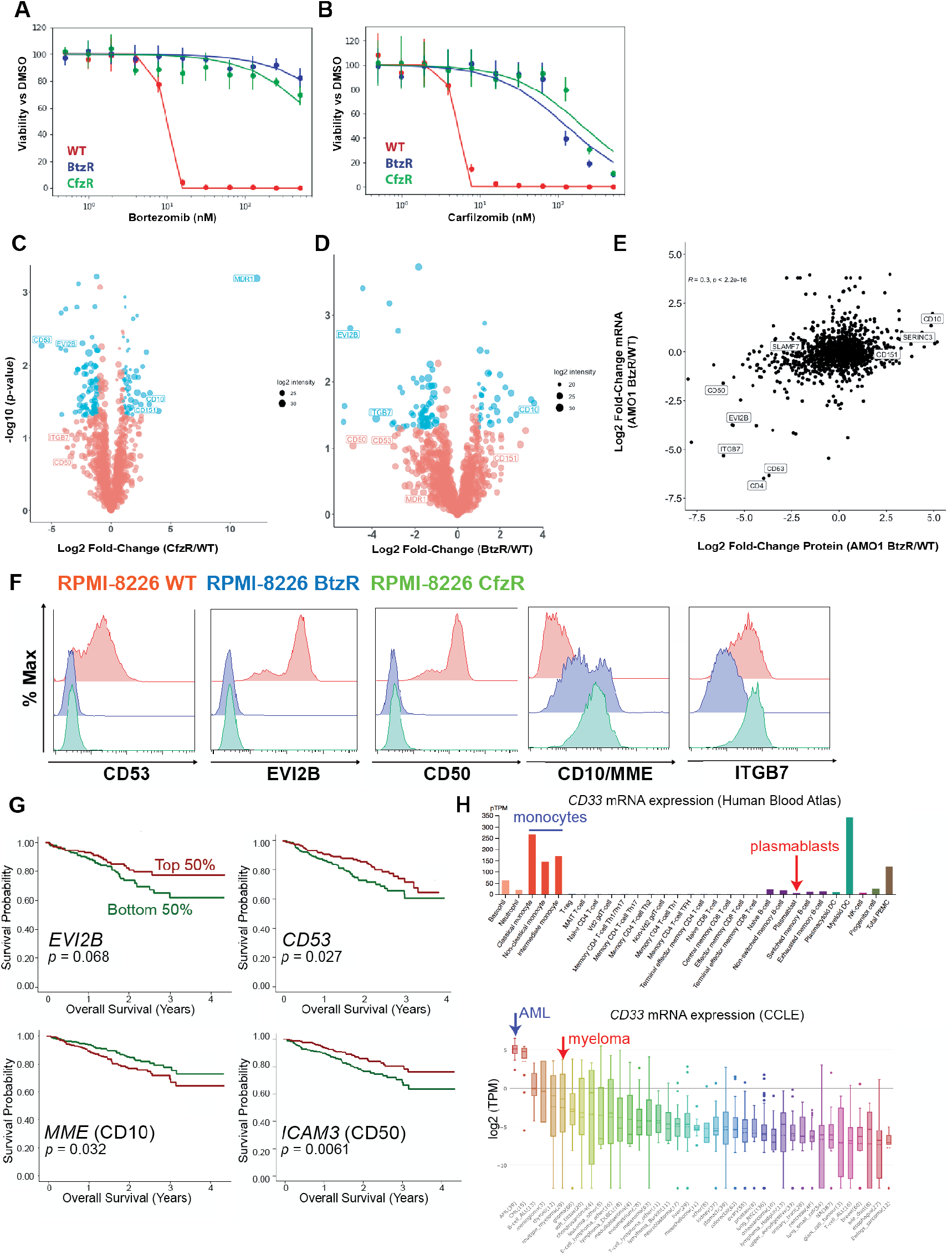
Additional characterization of surface biomarkers of myeloma drug resistance. **A-B.** Dose response curves for RPMI-8226 WT, BtzR, and CfzR evolved resistance lines treated with Btz (A) or Cfz (B) for 48 hours. **C-D.** Analogous to Fig. 3A, we display aggregate data of carfilzomib (C) or bortezomib (D) evolved resistant cell lines (CfzR and BtzR, respectively) versus their parental counterpart. Consistent with prior findings (35), we note that MDR1 is by far the most highly-upregulated surface protein in CfzR lines. Significantly-changed proteins noted in blue (log2-fold change >|1|; *p* < 0.05). **E.** Surface proteome-transcriptome comparison of BtzR AMO-1 vs. parental AMO-1 cells shows positive but relatively weak correlation (Pearson correlation and associated p-value shown). **F.** Flow cytometry confirms surface proteomic alterations in PI-resistant RPMI-8226 cells as also seen in AMO-1 cells (Fig. 3B). **G.** MMRF CoMMpass (Release IA10) overall survival data for tumor mRNA expression of noted putative PI-resistance biomarkers, separated by top (red) and bottom (green) half of expression. *p*-value by log-rank test. **H.** *CD33* is predicted to be expressed at far lower levels on plasmablasts (used as a proxy for plasma cells) than on normal myeloid lineage cells per data in the Human Blood Atlas (top), and myeloma cell lines express far less CD33 than AML cell lines in the CCLE (bottom).

**Supplementary Figure 5.**
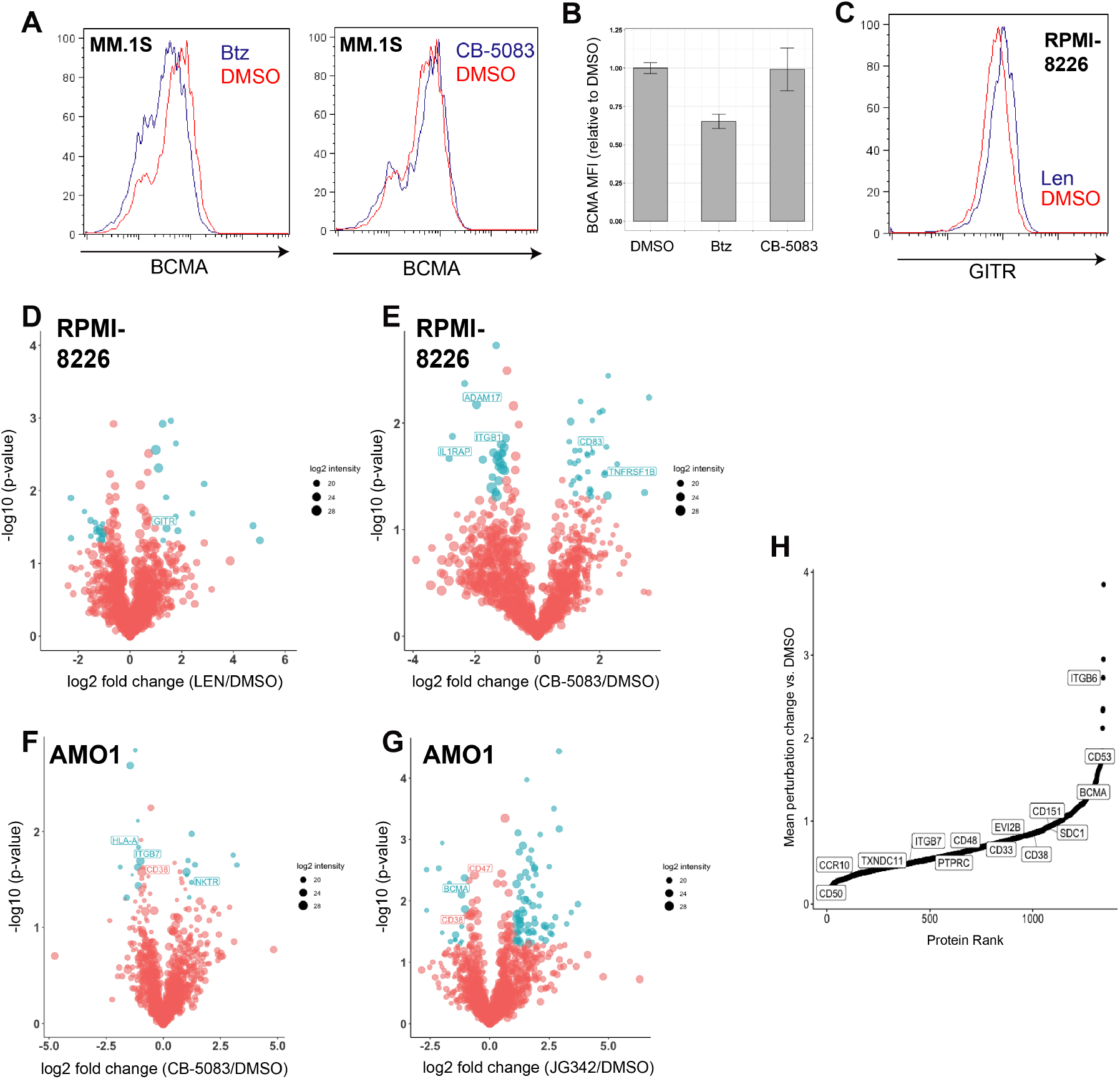
Probing myeloma surfaceome effects of acute drug treatment. **A.** MM.1S cells treated with 2.5 nM Btz for 48 hours show a decrease in surface BCMA as measured by flow cytometry, while treatment with 250nM CB-5083 for 48 hours shows no change. **B.** Quantification of results in (A) using median fluorescence intensity (*n*=2). **C.** RPMI-8226 cells treated with 25 μM Len show increase in GITR (*n* = 2). **D-G.** Profiling of RPMI-8226 cells treated with 50 μM Len for 48 hours (D), or 300 nM CB-5083 for 48 hours (E), and AMO1 cells treated with 250 nM CB-5083 for 48 hours (F), or 750 nM JG-342 for 48 hours (G) shows remodeling of the surface proteome (*n* = 3). **H.** Mean of protein absolute value change for 48 hour perturbations vs. DMSO for RPMI (Lenalidomide, Bortezomib, CB-5083) and AMO1 (Lenalidomide, CB-5083, JG342). Proteins on the right side of the distribution show the greatest variability across drug treatments while those on the left side show the least variability.

**Supplementary Figure 6.**
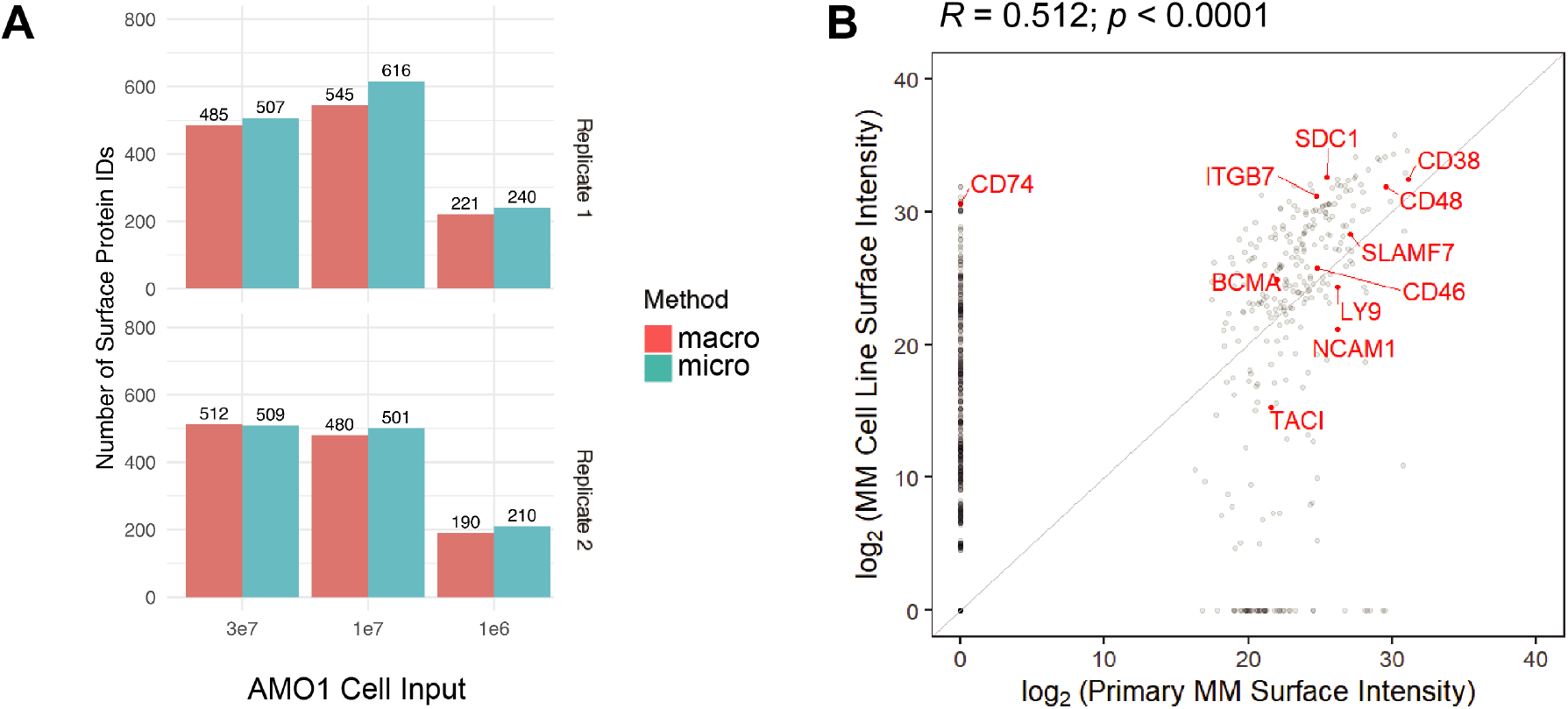
“Micro” method application to myeloma cell lines and primary sample. **A.** Surface protein identifications of the “micro” method applied to AMO1 myeloma cells, across a range of cell inputs, show less advantage over the standard “macro” method than initially found on RS4;11 B-ALL cells (Fig. 6B). **B.** Quantitative comparison of identified surface proteins (by LFQ intensity) in the assessed primary sample (*x*-axis) versus averaged over the four profiled myeloma cell lines *(y-* axis). We note very few proteins identified in the primary sample were not found in the assessed cell lines (i.e. few points with quantified LFQ intensity in the primary sample but 0 in cell lines). Many more proteins were identified in cell lines but not in the primary sample; however, this finding cannot determine whether these proteins are not truly expressed in the primary sample at all or were just unquantified due to lower overall proteomic coverage. Pearson correlation *R* value reported.

**Supplementary Table 1.** Scoring rubric to determine potential myeloma surface antigens for immunotherapeutic targeting. Assigned points based on perceived subjective importance to success of an immunotherapeutic strategy versus a specific antigen. Maximum possible score = 19. *Aggregate dataset includes cell surface capture proteomics on wild-type (RPMI-8226, AMO-1, L363, and KMS12-PE cells. **Non-hematopoietic cell types include all except “multiple_myeloma”, “B-cell_lymphoma_other”, “lymphoma_Burkitt”, “lymphoma_DLBCL”, “B-cell_ALL”, “T-cell_ALL”, “leukemia_other”, “T-cell_lymphoma_other”, “AML”, “CML”, “lymphoma_Hodgkin”. ***plasmablasts are used as a proxy for plasma cells as long-lived plasma cells are not analyzed as part of this dataset ****non-hematopoietic tissues include all except “Cells-EBV-transformed lymphocytes”, “spleen”, and “whole blood”.

| Dataset | 5 points | 4 points | 3 points | 2 points | 1 point | 0 points |
| --- | --- | --- | --- | --- | --- | --- |
| **Myeloma surface proteomics (this study)** |  |  | Identified in aggregate proteomic dataset* |  |  | Not identified in aggregate proteomic dataset |
| **COMPARTMENTS (subcellular localization prediction)** |  |  | Highest-confidence prediction at plasma membrane |  | Low-confidence prediction of plasma membrane localization | No predicted localization to plasma membrane |
| **Cancer Cell Line Encyclopedia** | Highest expression in myeloma cells, with >16-fold (4-fold on log2 scale) greater average expression than average in non-hematopoietic tumor types** |  | Highest expression in myeloma cells, but not meeting >16-fold threshold | Myeloma cell lines within top 5 most highly-expressing tumor cell types | Expressed in myeloma cells but not highly compared to other tumor types | Not expressed in myeloma cells |
| **Human Blood Atlas (Monaco et al. dataset)** |  | At least 10-fold higher expression in plasmablasts*** than any NK cells, T-cell, or myeloid cell type |  | Highest expression in plasmablasts but not meeting >10-fold criteria |  | Expressed but no enrichment in plasmablasts vs. other hematopoietic cells |
| **Genotype Tissue Expression (GTEx) project** |  | Average TPM in non-hematopoietic tissues**** <5 |  | Average TPM in non-hematopoietic tissues <50 |  | Average TPM in non-hematopoietic tissues >50 |

## SUPPLEMENTARY DATASET LEGENDS

**Supplementary Dataset 1. Mass spectrometry on Myeloma, B-lymphoblast, and Leukemia cell lines.**

**Supplementary Dataset 2. Output of ranking metrics for myeloma immunotherapy targets**

**Supplementary Dataset 3. Mass spectrometry on Proteasome-inhibitor resistant and WT Myeloma cell lines.**

**Supplementary Dataset 4. RNA sequencing on Proteasome-inhibitor resistant and WT Myeloma cell lines.**

**Supplementary Dataset 5. Mass spectrometry on Myeloma cell lines treated with small molecule inhibitors.**

**Supplementary Dataset 6. Mass spectrometry on primary patient sample using “micro” method.**

**Supplementary Dataset 7. Mass Spectrometry on Lenalidomide resistant and WT MM cell lines.**

**Supplementary Dataset 8. Membrane Protein lists use for filtering and data analysis.**

